# Single-cell multi-ancestry regulatory map of systemic lupus erythematosus

**DOI:** 10.64898/2026.09.18.752560

**Authors:** Haerin Jang, Catherine Sutherland, Wanseon Lee, Niek de Klein, Tarran S. Rupall, Bess L. Chau, Eliora Violain Buyamin, Norzawani Buang, Magdalena West, Chris Wincup, Katie L. Burnham, Emily R. Holzinger, Sarah Middleton, Frank Nestle, Emanuele de Rinaldis, Matthew C. Pickering, Marina Botto, Carla P. Jones, Timothy J. Vyse, James E. Peters, Gosia Trynka, Emma E. Davenport

**Author notes:** **Corresponding author:** Emma Davenport, Wellcome Sanger Institute, Wellcome Trust Genome Campus, Hinxton, United Kingdom, CB10 1SA.

## Abstract

Integrating genomic and single-cell transcriptomic profiles from patient cohorts can uncover regulatory effects underlying genetic associations with disease. Here we present SLEmap, a multi-ancestry single-cell expression quantitative trait locus (sc-eQTL) map generated from peripheral blood mononuclear cells from 281 patients with systemic lupus erythematosus (SLE). We identified 18,608 independent eQTLs in 5,656 genes, of which 149 signals in 66 genes colocalized with SLE GWAS loci. Nearly half of these colocalizations were detected exclusively through cell type-level analyses, highlighting the importance of cellular context for interpreting disease-associated genetic variation. Multi-ancestry data identified regulatory signals robust across diverse ancestral backgrounds and improved signal resolution. Most colocalizations were undetectable in sc-eQTLs from a healthy cohort (OneK1K), with novel SLE colocalizations highlighting disease-relevant regulatory mechanisms across distinct cell types, including the NF-κB pathway. These findings demonstrate the value of disease-specific, multi-ancestry single-cell regulatory maps for resolving the genes and cellular mechanisms underlying disease-associated genetic variation.

## Introduction

Most variants identified by genome-wide association studies (GWAS) as risk factors for complex diseases reside in non-coding regions, and their functional interpretation depends on connecting them to the genes and cell types through which they act. Expression quantitative trait loci (eQTL) mapping with bulk RNA sequencing (RNA-seq) has been instrumental in this effort, but averaging across cell types limits the resolution with which regulatory mechanisms can be identified. Studies in sorted immune cell populations^1–3^ and, more recently, at single cell resolution^4–7^ have shown that a large fraction of regulatory variants have effects restricted to specific cell types or states and suggested that these cell type-specific effects may colocalize more frequently with disease-associated loci. Large-scale single cell-eQTL (sc-eQTL) efforts, including OneK1K^4^, have begun to characterize this regulatory landscape systematically across immune cell types in healthy individuals. However, whether healthy cohorts fully capture disease-relevant regulatory variation remains an important open question.

Systemic lupus erythematosus (SLE) is a systemic autoimmune disease with over 330 associated GWAS loci^8^, the majority of which are non-coding and presumed to act through gene regulation. Multiple immune cell types are implicated in SLE pathogenesis, including both innate and adaptive immune cells, and prior work has demonstrated that regulatory effects at SLE loci are often cell type-specific^9^. Perez et al. provided early evidence for cell type-specific regulatory effects in SLE, profiling peripheral blood mononuclear cells (PBMCs) from 162 cases across eight immune cell types and colocalizing six genes with SLE GWAS loci. Yet, the majority of risk loci remain functionally uncharacterized, highlighting the need for greater cell type resolution, larger sample size, and broader patient population representation.

To address this, we present SLEmap, a multi-ancestry sc-eQTL map generated from 281 female patients with SLE, with matched whole-genome sequencing (WGS) and single-cell RNA sequencing (scRNA-seq) data. We identify 18,608 conditionally independent eQTLs in 5,656 unique genes and show that cell type-level mapping recovers regulatory signals missed in aggregate analyses. Colocalization with a large multi-ancestry SLE GWAS identifies 149 colocalizations across 66 genes, of which 23 are previously unreported in bulk or single-cell studies, and the majority are not detectable in a large healthy cohort (OneK1K^4^), underscoring the value of a disease-specific, multi-ancestry sc-eQTL map for dissecting the genetic architecture of SLE.

## Results

### SLEmap: a multi-ancestry single-cell eQTL map of SLE

We assembled a multi-ancestry cohort of SLE patients to resolve cell type-specific genetic regulation of gene expression in a disease context (**Fig 1A, B**). We first performed multi-omic single-cell RNA-seq on a total of 663,433 PBMCs from 288 female SLE patients and 10 healthy controls. Integrated analysis of surface protein expression (CITE-seq), RNA markers, and immune receptor (BCR/TCR) data enabled annotation of 26 immune cell types aligned with flow cytometry-based immunophenotyping (**Fig. 1C, D; Supplementary Figs. 1, 2**). For cell type-resolved eQTL mapping in the SLE disease context, we used 281 SLE patients with matched single cell and WGS data (SLEmap cohort, **Fig. 1A**). A defining feature of SLEmap is its multi-ancestry composition (**Fig. 1B**), with the three largest groups having genetically inferred African (AFR, n=102), European (EUR, n=81), and South Asian (SAS, n=62) ancestries, enabling identification of regulatory effects generalizable across SLE populations.

**Figure 1.**
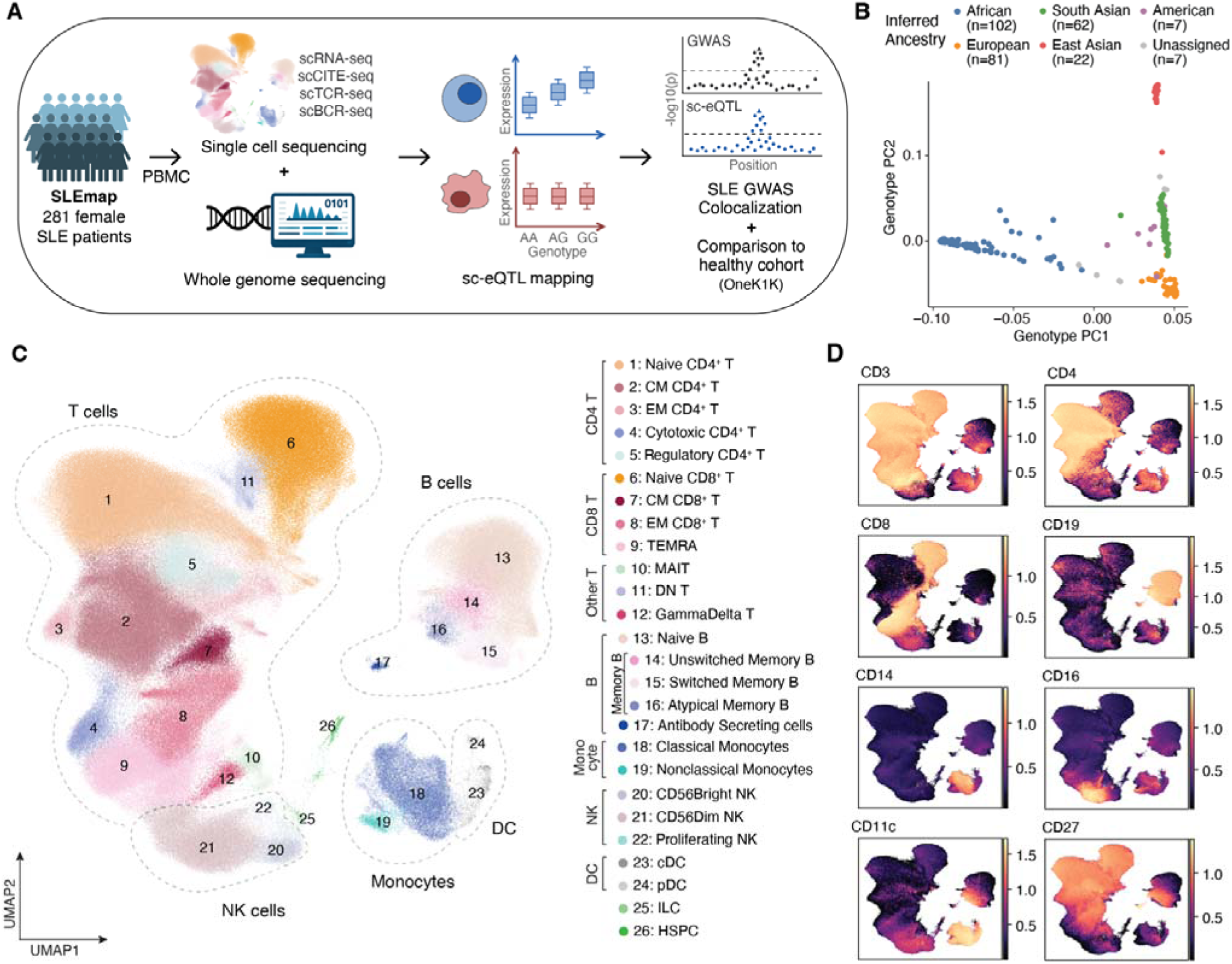
SLE sc-eQTL mapping study design and multimodal immune cell annotation. **(A)** Overview of SLEmap cohort and eQTL mapping study design. Matched single-cell and WGS data was available from 281 female SLE patients. Cell type-resolved sc-eQTL mapping was conducted to perform colocalization with SLE GWAS loci. **(B)** Cohort ancestry composition, inferred from genotype data. **(C)** UMAP of 663,433 PBMCs from 288 SLE patients and 10 healthy controls annotated into 26 immune cell types, generated and clustered using integrated CITE-seq and scRNA-seq data. **(D)** Surface protein expression of canonical cell type markers (orange=high expression, purple=low expression). CM=central memory; EM=effector memory; TEMRA=terminally differentiated effector memory RA cells; DN=double negative; MAIT=mucosal-associated invariant T; NK=natural killer; cDC=classical dendritic cell; pDC=plasmacytoid dendritic cells; ILC=innate lymphoid cells; HSPC=hematopoietic stem and progenitor cells

### Cell type-level sc-eQTL mapping reveals regulatory signals masked in bulk analyses

We mapped conditionally independent cis-eQTLs across 15 immune cell types with sufficient cell numbers and individuals. Gene expression was quantified using a mean pseudobulk approach, restricted to cell types with at least 100 patients contributing a minimum of 20 cells each to ensure reliable expression estimates. eQTLs were also mapped in all cells combined (all-PBMC, 622,452 cells), where expression profiles were derived from all cells regardless of cell type, to maximize eQTL discovery and characterize how cell type-level regulatory signals differ from those detected in aggregate. In total, 11,899 conditionally independent eQTLs corresponding to 3,803 unique eGenes were identified across 15 cell types, and 6,709 conditionally independent eQTLs in 4,502 unique eGenes were identified in all-PBMC (**Fig. 2A, Supplementary Table 1**). Overall, one or more eQTLs were identified in 5,656 genes. The number of eQTLs varied considerably across cell types, reflecting differences in statistical power driven by the number of cells and individuals per cell type, both of which correlated with the proportion of significant eGenes detected (**Supplementary Fig. 3**). Accordingly, the all-PBMC analysis, which included the largest number of cells and individuals, identified the greatest number of eQTLs.

**Figure 2.**
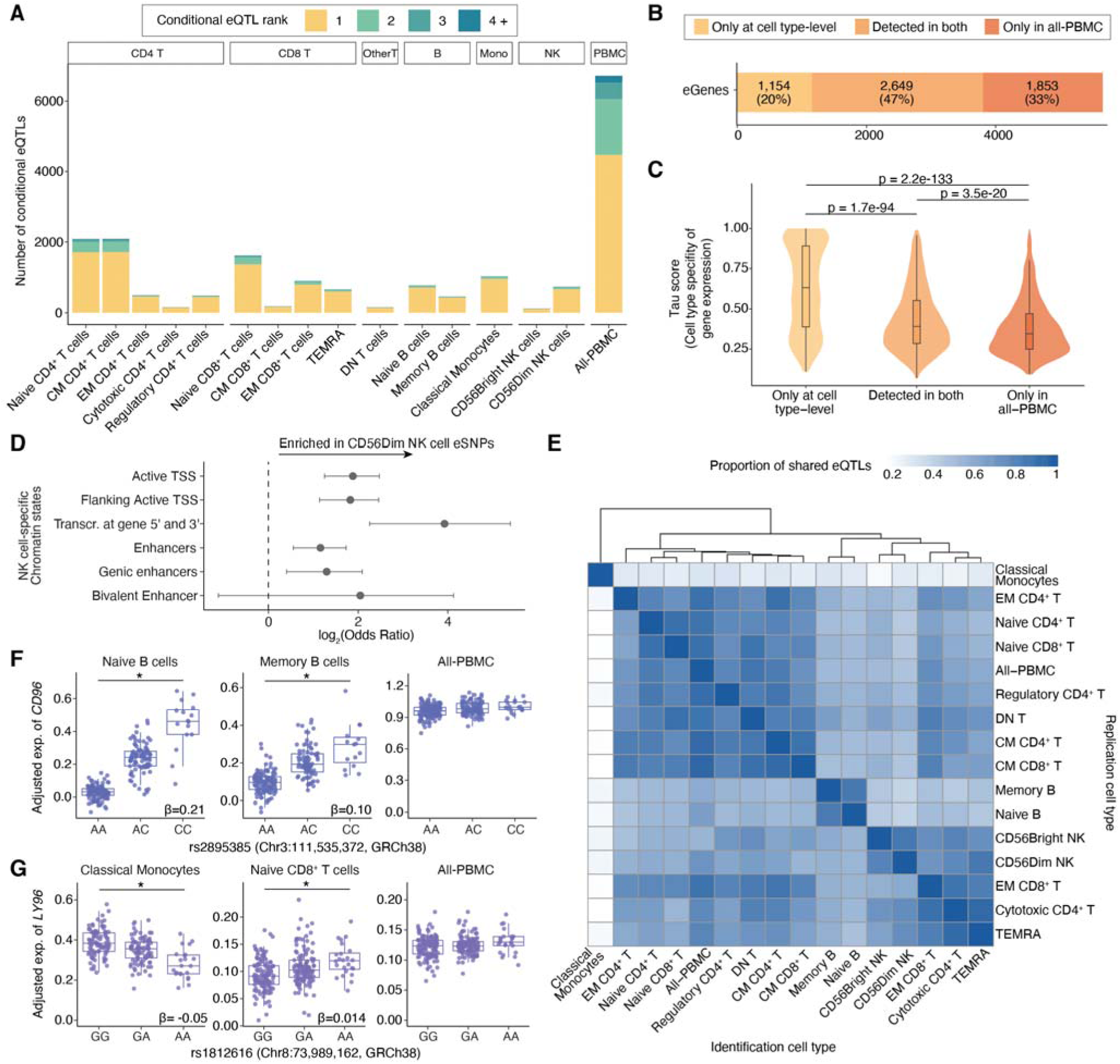
Conditionally independent sc-eQTL mapping and comparison to eQTLs mapped with all-PBMC. **(A)** Number of conditionally independent eQTLs identified by cell type. All-PBMC denotes eQTL mapping performed using pseudobulk expression from all cells in the single cell dataset combined. **(B)** Unique eGenes identified from cell type-level eQTL mapping compared to all-PBMC. For cell type-level results, eGenes detected in multiple cell types were counted once. **(C)** Tau score of eGenes detected in all-PBMC and cell type-level eQTL mapping. Tau score measures cell type specificity, with 0 meaning broadly expressed across multiple cell types, while 1 means cell type specific expression. Box plots indicate the interquartile range (box), median (center line), and minimum/maximum values excluding outliers (whiskers). P-values were calculated using the Wilcoxon rank sum test, indicating the significance of differences between groups. **(D)** Enrichment of NK cell-specific chromatin states among eSNPs identified in CD56DimNK cells against all-PBMC. Only significant enrichments are shown; error bars indicate 95% confidence intervals. Results for all immune cell types with matching primary peripheral blood cell annotations are shown in **Supplementary Fig. 5**. **(E)** Pairwise eQTL sharing between cell types, assessed by mashr. Color indicates the proportion of eQTLs significant in the identification cell type that are shared (significant with consistent effect size) in the replication cell type. **(F)** Example of an eQTL shared within a lineage but not in all-PBMC (*CD96*-rs2895385). * = significant eQTL. **(G)** Example of an eQTL with opposite effects between cell types (*LY96*-rs1812616) and not detected in all-PBMC. Gene expression levels are adjusted for covariates used in the eQTL model. TSS=Transcription start site

Cell type-level mapping revealed 1,154 unique eGenes not found in the all-PBMC analysis (**Fig. 2B**), largely representing genes expressed in a cell type-specific manner whose signals are likely masked when cell types are pooled (**Fig. 2C**) and whose pseudobulked expression may be affected by variation in differential cell proportions across individuals. Conversely, eGenes identified exclusively in the all-PBMC analysis had lower cell type specificity (**Fig. 2C**) and lower overall expression compared to those identified in both analyses (**Supplementary Fig. 4**). This suggests the signals detected only in all-PBMCs reflect either genes with broad but low-level expression across most immune cells, or cell type-specific signals underpowered at the individual cell type level. Furthermore, compared to those identified in all-PBMC, cell type-level eSNPs were more strongly enriched in cell type-specific regulatory chromatin states, particularly at active transcription start sites and enhancer elements (**Fig. 2D, Supplementary Fig. 5**), indicating that the two approaches capture not only different eGenes but distinct classes of regulatory variation.

To further characterize eQTL sharing and specificity across cell types, we assessed pairwise sharing across cell type pairs using mashr^10^, which models effect size heterogeneity while accounting for differences in statistical power between cell types (**Fig. 2E**). Shared effects were more frequent within related immune lineages, such as naive and memory B cells, and among functionally related cell types, such as natural killer (NK) cells, terminally differentiated effector memory T (TEMRA) cells, and cytotoxic CD4^+^ T cells, which share cytotoxic properties. Notably, all-PBMC clustered with T cell subtypes, consistent with T cells comprising the largest proportion of the single-cell dataset (77%) and therefore having the greatest influence on the combined pseudobulk profiles. As the only myeloid lineage cell type included in the analysis, classical monocytes were the most distinct, showing the lowest proportion of shared eQTLs with any other cell type.

While mashr captures broad patterns of eQTL sharing across cell types, we next sought to determine whether individual eQTL signals were shared between specific cell type pairs. We therefore performed fine-mapping and colocalization across cell types (**Supplementary Fig. 6**). This identified cases where a genomic region harbored shared and distinct underlying signals. For example, at the *CD96* locus, there was a shared signal between naive and memory B cells not found in all-PBMC (**Fig 2F**). This signal was distinct from a nearby signal shared across CD4^+^ T cell subsets (**Supplementary Fig. 7**). A subset of signals showed opposite effects on gene expression between cell types (**Supplementary Table. 2**) and were often undetectable in all-PBMC eQTLs as illustrated by the *LY96* locus (**Fig. 2G**). Together, these results reinforce that cell type-level mapping captures a distinct and biologically meaningful layer of regulatory variation that is obscured in bulk analyses.

### sc-eQTL colocalization with SLE GWAS identifies candidate genes and supports cross-ancestry generalisability

Having demonstrated that eQTL mapping at the cell type-level recovers regulatory signals missed in aggregate analyses, we next asked whether this resolution translates into improved interpretation of SLE GWAS loci. Conditionally independent sc-eQTLs were colocalized with the multi-ancestry SLE GWAS meta-analysis by Khunsriraksakul et al.^11^ using coloc^12^, testing GWAS loci reaching genome-wide suggestive significance (p < 1e-5). Of 197 GWAS loci tested, 53 colocalized with at least one eQTL (PP.H4 > 0.8), implicating 66 candidate genes through 149 total colocalizations across loci, genes, and cell types (**Fig. 3, Supplementary Table 3**). To assess novelty, colocalizations were compared against all known eQTL-SLE GWAS colocalizations in the Open Targets Platform, including results from both bulk and sc-eQTL studies. Of the 66 colocalized eGenes, 23 (38%) were not previously reported.

**Figure 3.**
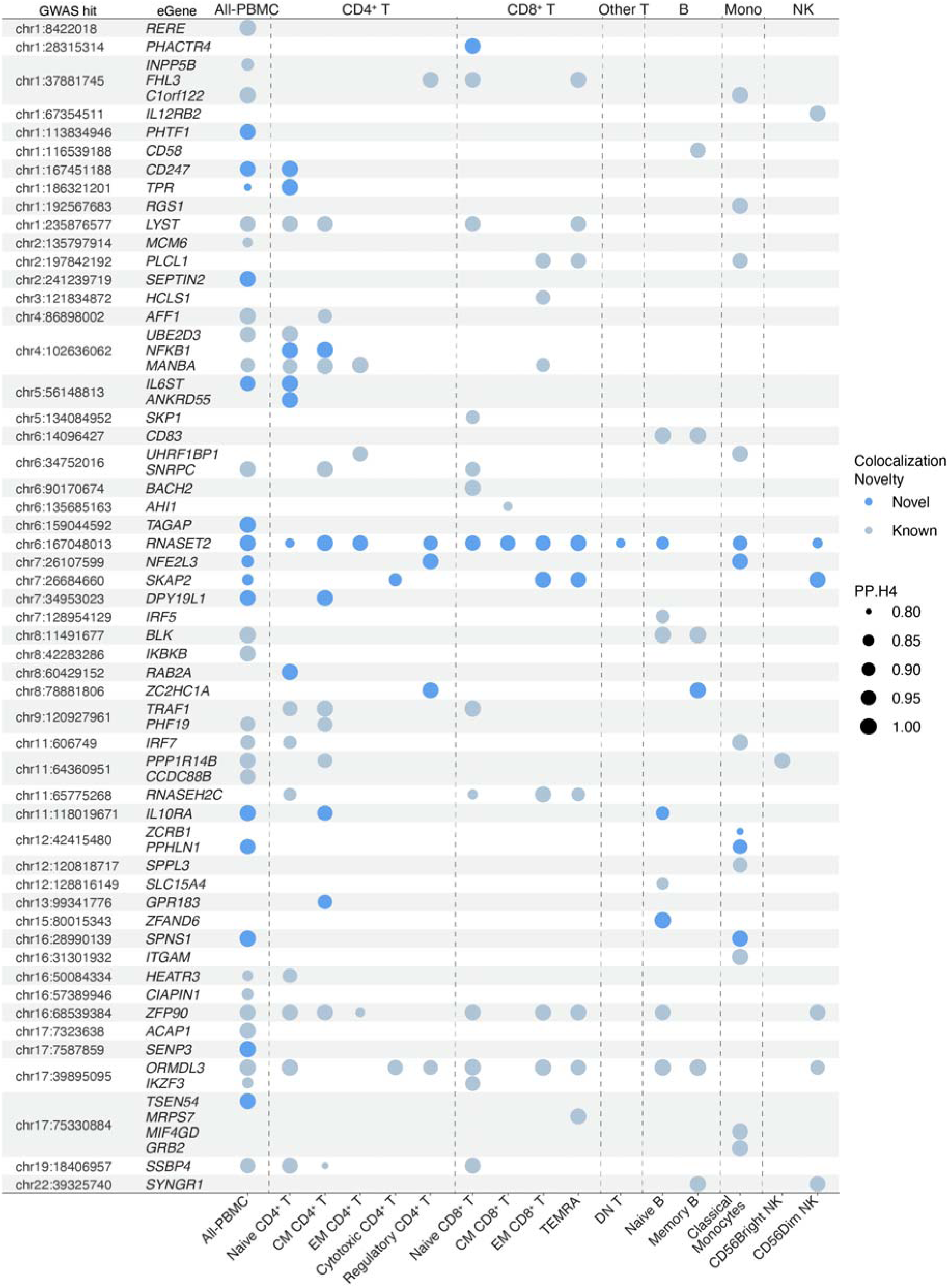
SLE GWAS loci colocalizing with sc-eQTLs. The size of each dot represents the PP.H4 for colocalization. Novelty was assessed by comparing to all known colocalizations (PP.H4 > 0.8) between eQTL (single cell or bulk) and SLE GWAS studies reported in the Open Targets database. GWAS loci ordered by chromosome and position (GRCh38). Each shaded row shows all eGenes colocalizing with the same GWAS loci.

Given the multi-ancestry composition of SLEmap, we next asked whether these regulatory effects were shared across ancestry groups. Many regulatory effects are reported to be conserved across ancestry groups, though differences in allele frequency and study power can determine whether such effects are detectable in any given population^13^. To assess sharing of eQTL effects across ancestries, sc-eQTLs were mapped genome-wide within each ancestry group and effect sizes of colocalized eQTLs were compared across AFR, EUR, and SAS. Effect sizes were broadly correlated across all three groups (Pearson’s r = 0.88-0.89, p < 0.05 for all pairwise comparisons; **Fig. 4A**), indicating that the majority of detected regulatory effects are shared.

**Figure 4.**
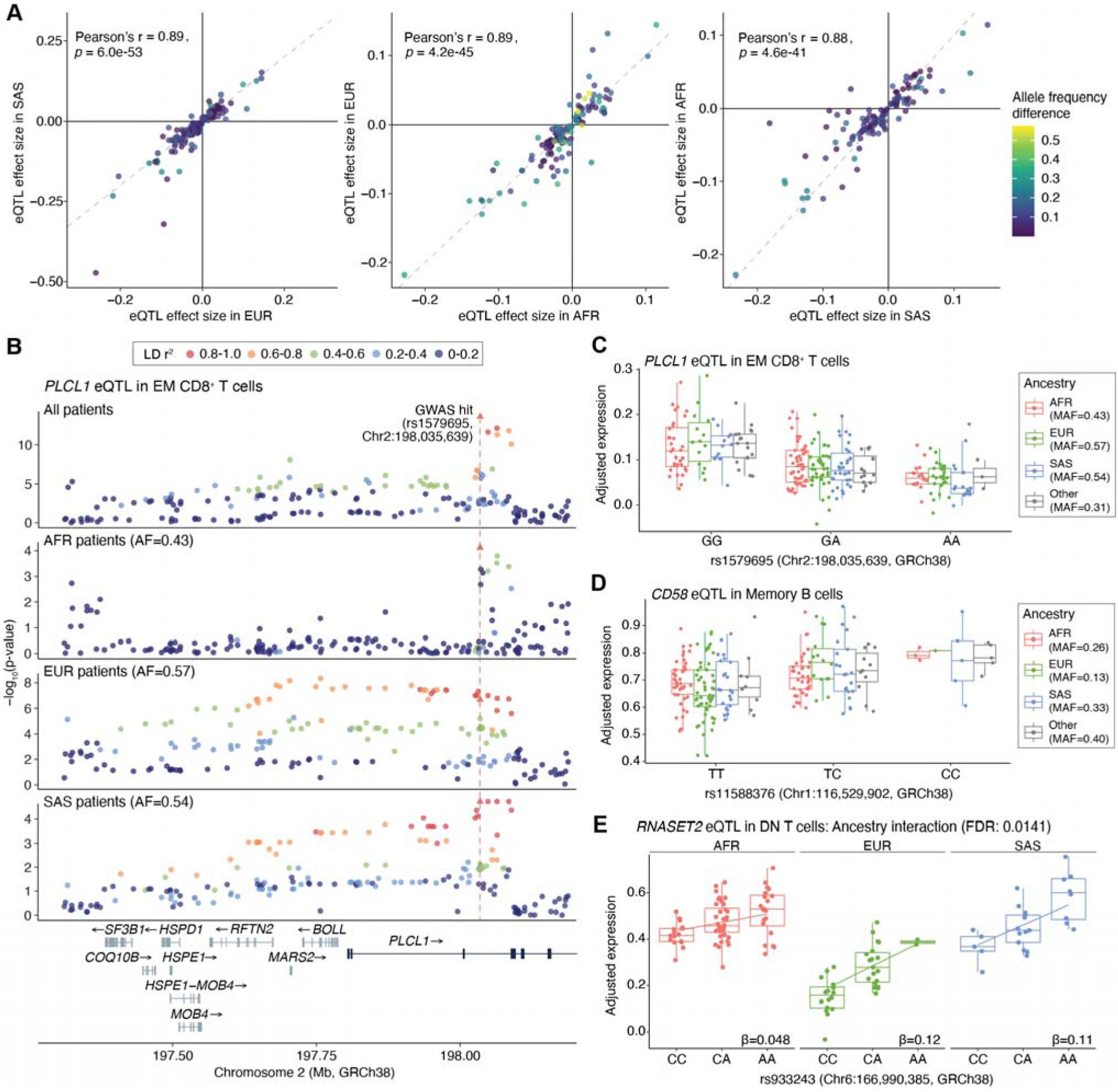
Multi-ancestry eQTL analysis reveals shared and ancestry-specific genetic effects. **(A)** Correlation of eQTL effect sizes for colocalized signals across ancestry groups (AFR, EUR, SAS), with points colored by inter-ancestry allele frequency difference. Only eSNPs with > 0.05 MAF in each ancestry group are shown. **(B)** eQTL regional plot of *PLCL1* eQTL in EM CD8^+^ T cells, illustrating how multi-ancestry data refines signal resolution. Points are colored by LD r² to the GWAS hit calculated within each ancestry group. Gene tracks generated with locuszoomr. **(C)** Box plot of the *PLCL1* eQTL colored by ancestry. **(D)** Box plot of a *CD58* eQTL in memory B cells, where the effect allele has low allele frequency in Europeans yet shows a consistent effect across ancestries. **(E)** Box plot of an ancestry-interaction eQTL in *RNASET2* in DN T cells.

The multi-ancestry design did however improve resolution of the regulatory architecture at several loci. At the *PLCL1* locus in EM CD8^+^ T cells, inclusion of multiple ancestry groups refined the underlying association pattern, pointing towards a more precise candidate regulatory variant driving the shared effect (**Fig. 4B, C**). In EUR, multiple variants were in high linkage disequilibrium (LD) with the lead variant, producing a broad association peak. In contrast, the short LD block observed in AFR reduced this correlated set, allowing the shared association to be localized to a smaller number of candidate variants when analyzed jointly. The multi-ancestry design also enabled detection of regulatory effects at variants that would be underpowered or untestable in certain single-ancestry studies. At the *CD58* locus in memory B cells, the colocalized variant is relatively rare in EUR (minor allele frequency (MAF) 0.13 compared to 0.26 in AFR and 0.33 in SAS) yet shows a consistent effect across all three ancestry groups (**Fig. 4D**).

To assess whether any colocalizations showed evidence of ancestry-specific regulation, we tested for ancestry interaction-eQTLs among the 74/149 colocalizations that passed stringent filters (see **Methods**). A single significant interaction was identified: *RNASET2* in DN T cells (FDR = 0.014, **Fig. 4E**). The SLE risk allele (C) at rs933243 (chr6:166,990,385) was associated with reduced *RNASET2* expression. The eQTL effect was stronger in EUR and SAS than in AFR. In addition, baseline *RNASET2* expression was lower in EUR than in AFR and SAS, indicating that ancestry may influence *RNASET2* expression both independently of genotype and by modifying the magnitude of the risk-allele effect. Thus, EUR individuals carrying the risk allele have both lower baseline *RNASET2* expression and a stronger reduction associated with the risk allele. Notably, *RNASET2* colocalized across 12/15 cell types, yet the ancestry interaction was significant only in DN T cells, suggesting that ancestry-specific regulatory effects may themselves be cell type-specific. Though current sample sizes within individual ancestry groups limit the power to detect such interactions, larger multi-ancestry cohorts hold promise for systematically uncovering the prevalence of ancestry-specific regulatory effects in SLE.

### The majority of SLE-relevant colocalizations are not detected in a large healthy cohort

To better understand the contribution of disease-specific regulatory context in the detection of colocalizations, we conducted a more thorough assessment of how these colocalizations compared to a large single cell eQTL cohort generated on healthy individuals (OneK1K^4^, a European ancestry cohort). To make the dataset comparable and avoid confounding by sex, we subsetted the dataset to female individuals (n = 565) and annotated cell types to align with SLEmap (**Fig. 5A; Supplementary Figs. 8, 9**). Identical eQTL mapping and colocalization methods were applied across both datasets, with 13 overlapping cell types and all-PBMC tested in both (**Supplementary Table 4**).

**Figure 5.**
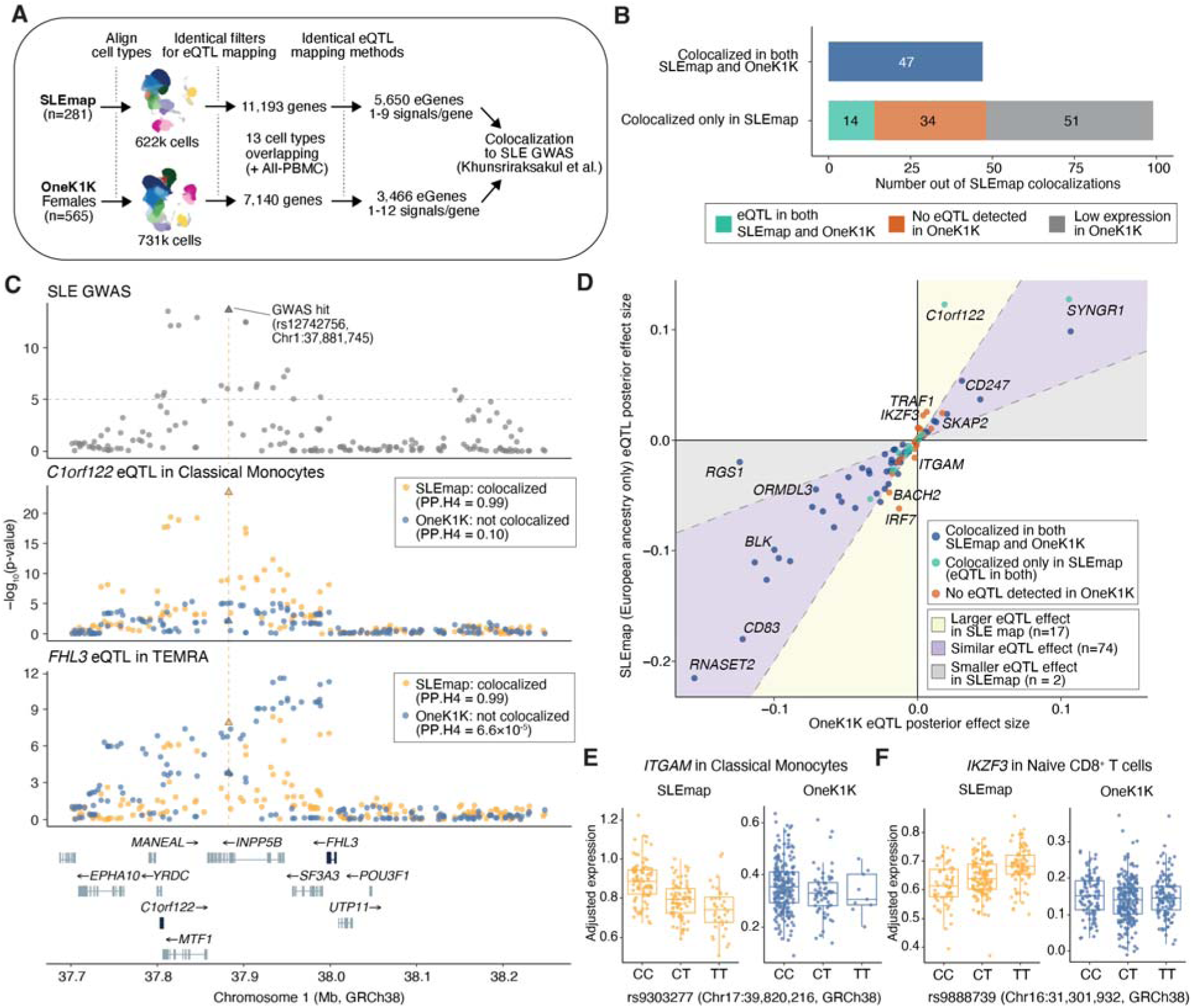
Comparison of SLEmap eQTL colocalizations with the OneK1K healthy cohort. **(A)** Schematic of the comparison pipeline. Cell types were harmonized across datasets using shared RNA markers. Identical filters were applied for eQTL mapping (genes expressed in >5% of cells and >10% of individuals, >20 cells per individual per cell type, >100 qualifying individuals per cell type). Gene counts were restricted to overlapping cell types. **(B)** Number of colocalized GWAS locus-eGene-cell type combinations in SLEmap replicated in OneK1K. Colocalizations in cell types absent from OneK1K are excluded. **(C)** Locus plot of the 1p34.3 SLE risk locus (upper panel). SLE risk colocalizes with *C1orf122* eQTL in classical monocytes and *FHL3* eQTL in TEMRA cells in SLEmap but not OneK1K. Gene tracks generated with locuszoomr. **(D)** Shared and distinct eQTL effect magnitudes in colocalized signals between OneK1K (European ancestry cohort) and the European-only SLEmap subset. Posterior effect sizes calculated using mashr. **(E)** Box plot of the *ITGAM* eQTL in classical monocytes, significant eQTL detected in SLEmap but not OneK1K. **(F)** Box plot of the *IKZF3* eQTL in naive CD8^+^ T cells, significant eQTL detected in SLEmap but not OneK1K.

Out of the 146 SLEmap colocalizations detected in cell types tested in OneK1K (13 cell types, 66 genes), 47 (32%) were also detected in OneK1K (**Fig. 5B, Supplementary Table 3**). Among the remaining colocalizations, 14 (10%) had a detectable eQTL in OneK1K for the same cell type that did not colocalize, 34 (23%) had no detectable eQTL for the gene in OneK1K, and 51 (35%) could not be tested due to insufficient gene expression.

The majority of SLEmap colocalizations were therefore not detectable in OneK1K, likely reflecting a combination of technical, study design, ancestry, and disease-specific differences. The large number of genes with insufficient expression in OneK1K may partly reflect disease-specific expression but could also result from differences in sequencing depth (mean UMI per cell: 5,075 in SLEmap vs. 3,476 in OneK1K) and cells profiled per individual (mean: 2,215 versus 1,294). Differences in the underlying heterogeneity within the cell types defined in SLEmap and transferred to OneK1K could also reduce power to detect eQTL in the latter. Conversely, OneK1K included more individuals than SLEmap (565 versus 281), providing greater power to detect eQTLs and a larger total number of cells. Given our interest in colocalizations that may reflect disease context, we focused on cases where sufficient expression and statistical power supported eQTL detection in both cohorts, but colocalization was observed only in SLEmap.

The 1p34.3 locus illustrates two mechanistically distinct reasons why colocalizations may be SLEmap-specific even when there is a significant eQTL detected in OneK1K (**Fig. 5C**). For C1orf122 in classical monocytes, the stronger eQTL significance in SLEmap may explain the stronger colocalization and could reflect context-specific regulation. For *FHL3* in TEMRA cells, eQTL significance was similar between datasets but the association patterns differed markedly; as this difference persisted in the European-only SLEmap subset (**Supplementary Fig.10**), ancestry composition was unlikely to be the primary driver raising the possibility that disease state alters the local regulatory architecture at this locus.

To enable a more direct comparison of disease status while adjusting for the influence of ancestry and potential differences in statistical power between datasets, eQTL effect sizes for SNVs with the highest PP.H4 in each colocalization in SLEmap were compared between European-ancestry individuals from SLEmap and OneK1K using mashr^10^ (**Fig. 5D, Supplementary Fig. 11**). Most colocalizations showed shared effects between SLEmap and OneK1K (posterior effect size ratio within a factor of two, 74/93 colocalizations). However, a subset (17/93 colocalizations), including those of key SLE-associated genes such as *IRF7* in classical monocytes and *BACH2* in naïve CD8 T cells, showed substantially larger posterior effect sizes in SLEmap. Additionally, some colocalized eQTLs were undetectable in OneK1K, including *ITGAM* in classical monocytes and *IKZF3* in naive CD8^+^ T cells (**Fig. 5E, F**). Ancestry differences did not account for these discrepancies (**Supplementary Fig. 12**), pointing to disease state as a likely driver.

### sc-eQTL colocalization with SLE GWAS identifies cell type-specific disease mechanisms

Among genes colocalizing with SLE GWAS loci in SLEmap, 46% were identified only through cell type-level eQTL mapping, compared with 14% identified exclusively through all-PBMC mapping (**Fig. 6A, Supplementary Fig. 13**), highlighting the value of sc-eQTL mapping in considering cellular context when interpreting genetic disease risk. For example, *IL6ST* is broadly expressed across immune cells, but our data specifically implicates its genetic regulation in naive CD4^+^ T cells, where it is known to integrate IL-6 and IL-21 signaling to promote Th17 and Tfh differentiation over Tregs, leading to chronic inflammation in SLE^14–17^. Among the colocalized genes, two recurring themes were nucleic acid sensing and type I interferon production, and NF-κB inflammatory signaling, with different genes implicated across cell lineages (**Fig. 6B**). For instance, in naïve B cells, colocalizations at *IRF5* and *SLC15A4* were identified exclusively in this cell type and were absent from OneK1K, pointing to a disease-relevant, cell type-restricted regulatory effect. *SLC15A4* is required for downstream signaling from TLR7/9 sensing of nucleic acid-containing immune complexes, and *IRF5* acts as the direct downstream transcription factor (TF) driving type I interferon and pro-inflammatory cytokine production from this signal. Their co-identification reinforces the importance of this pathway in B cell-mediated immune response in disease. Notably, *SLC15A4* inhibition has been suggested as a therapeutic strategy in autoimmune conditions, lending translational relevance to this finding^18^.

**Figure 6.**
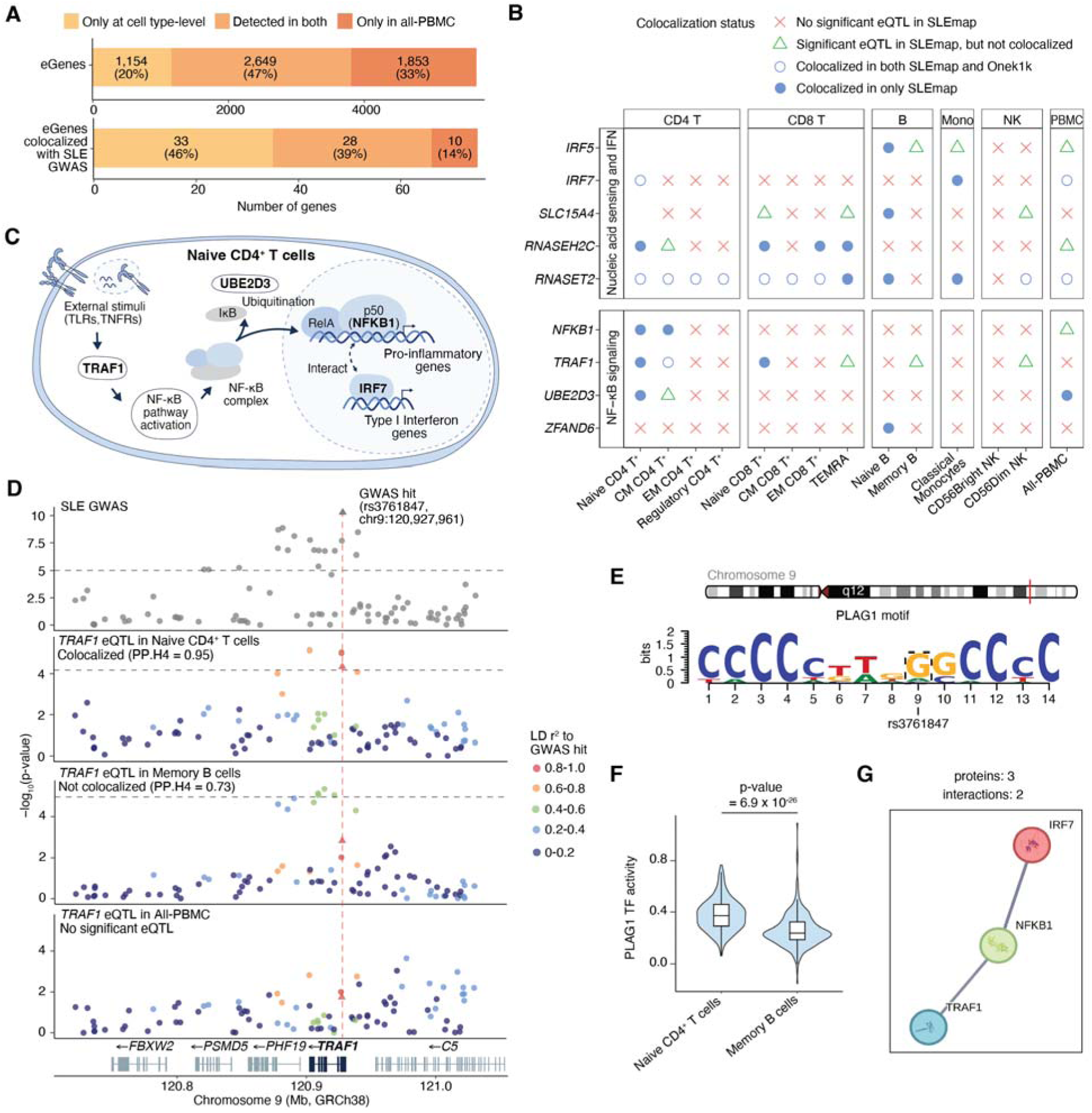
Colocalization of sc-eQTLs with SLE GWAS identifies cell type-specific regulatory mechanisms. **(A)** Number and proportion of genes identified by cell type-level and all-PBMC eQTL mapping that colocalize with SLE GWAS loci. **(B)** sc-eQTL status for colocalized genes related to ‘nucleic sensing and interferon’ and ‘NF-kB signaling’ across cell types. No symbol denotes those not tested for eQTLs due to insufficient expression (expressed in fewer than 5% of cells or fewer than 10% of donors). For genes with multiple independent sc-eQTLs, the sc-eQTL with the higher likelihood of colocalization was plotted. **(C)** Schematic of potential interaction between colocalized genes (*TRAF1, UBE2D3, NFKB1, IFR7*) in naïve CD4^+^ T cells leading to immune response in SLE. **(D)** Colocalization at 9q33.2 GWAS locus with *TRAF1* in naive CD4^+^ T cells showing cell type-specific colocalization. Points are colored by LD r^2^ to the GWAS hit. The dotted grey line indicates the suggestive genome-wide significance threshold (P = 1 × 10⁻) for GWAS and nominal p-value threshold for eQTL. **(E)** PLAG1 TF binding motif disruption by GWAS risk variant assessed using motifbreakR. **(F)** PLAG1 TF activity in naive CD4^+^ T cells and memory B cells measured using decoupleR. Box plots indicate the interquartile range (box), median (center line), and minimum/maximum values excluding outliers (whiskers). P-values were calculated using a two sample t-test to assess the difference between cell types. **(G)** Protein-protein interaction network of *IRF7, NFKB1*, and *TRAF1*. Network generated using STRINGdb. Nodes show 3D protein structure and edges indicate predicted functional associations.

In naïve CD4 T cells, three genes (*TRAF1*, *UBE2D3,* and *NFKB1)* which colocalized in SLEmap but not OneK1K, converge on the NF-κB signaling pathway (**Fig. 6B, C**). *TRAF1* functions as an adaptor downstream of TNF receptor and TLR signaling to activate NF-κB^19,20^, *UBE2D3* promotes IκBα ubiquitination and degradation enabling nuclear translocation of the NF-κB complex^21^, and *NFKB1* encodes p50, a core component of the NF-κB complex^22^. In particular, the colocalization of *TRAF1* illustrates how regulatory effects at the same locus can differ across immune populations (**Fig. 6D**). Although an eQTL of similar strength was observed at the same locus in memory B cells, this signal did not colocalize with either the GWAS or the naïve CD4 T cell eQTL, while in all-PBMC no eQTL was detected. This indicates that distinct regulatory variants drive *TRAF1* expression in each cell population, of which only those active in specific cell types are associated with SLE risk.

To investigate the functional basis of this cell type-specific effect in *TRAF1*, we further assessed whether the colocalized variant (rs3761847, chr9:120,927,961) disrupted TF binding motifs, identifying predicted disruption of PLAG1, TFAP2, TFAP2A, and TFAP2C binding sites. After filtering to TFs with detectable expression in PBMCs, only PLAG1 remained (**Fig. 6E**). Inferred PLAG1 TF activity was significantly higher in naïve CD4 T cells than in memory B cells (**Fig. 6F**), consistent with a mechanism by which the colocalized variant more strongly affects *TRAF1* expression through disruption of PLAG1 binding in this cell type.

Finally, we explored potential connectivity between colocalizing genes in naive CD4 T cells by querying the STRING database for high confidence interactions (>700). This suggested an interaction between *NFKB1*, *TRAF1*, and *IRF7* based on co-expression networks (**Fig. 6C, G**). *IRF7* operates within a parallel interferon regulatory pathway that cross-talks with NF-κB to coordinate cytokine production and antiviral responses^23^. The convergence of these genes within specific cell types illustrates how sc-eQTL results can move beyond individual associations to implicate shared immune signaling networks through which polygenic SLE risk may act and provides a framework for prioritizing mechanistic follow-up.

## Discussion

In this study, we present SLEmap, a multi-ancestry sc-eQTL map generated from patients with SLE through integration of matched single-cell transcriptomic and WGS data. Our analyses demonstrate that cell type-specific regulatory mapping in the context of disease improves interpretation of SLE risk loci, enabling more accurate assignment of genetic associations to the genes and immune cell populations through which they are likely to act. SLEmap identifies 66 colocalized genes, markedly expanding the proportion of SLE GWAS loci with a prioritized target gene and cell type compared to previous SLE-specific studies^24^, and also including several loci previously unreported in bulk transcriptomic studies or healthy cohorts.

Many of these colocalizations revealed strong cell type-specificity, with nearly half identifiable only through cell type-level mapping. This is consistent with growing evidence that risk variants of complex traits frequently exert context-dependent effects restricted to particular cell types rather than acting broadly across tissues^1,5,25,26^. By resolving the specific immune cell types in which these regulatory effects occur, cell type-level colocalization can help refine otherwise ambiguous GWAS signals into more mechanistically interpretable effector genes and generate hypotheses about their functional context. For example, colocalization with a *SKAP2* eQTL occurred only in cytotoxic cell types (CD56Dim NK cells, EM CD8^+^ T cells, TEMRA, and cytotoxic CD4^+^ T cells). As a signaling adaptor downstream of FcγRs and integrins, *SKAP2* regulates immune cell function and the adhesion programs required for tissue infiltration ^27^, suggesting a role in the migration of cytotoxic cells that may contribute to tissue inflammation in SLE. Similarly, *SPNS1* colocalizes exclusively in monocytes, where its function as a lysosomal transporter ^28^ may implicate it in the endolysosomal compartment through which TLR7 and TLR9 sense apoptotic nucleic acids and drive the type I interferon signature characteristic of SLE. Together, these findings demonstrate the utility of SLEmap as a map for linking SLE risk loci to the genes, cell types, and biological mechanisms through which they may contribute to disease and, hence informing future therapeutic studies.

Beyond individual loci, these colocalizations converge on shared immune pathways related to nucleic-acid sensing, interferon signaling, NF-kB signaling, and BCR activation, illustrating how eQTL-GWAS colocalization links disease-associated variants to the cellular pathways and immune populations underlying pathogenesis. In monocytes, colocalization at *IRF7*, *RNASET2* and *PPHLN1* points to dysregulated type I interferon and endolysosomal sensing pathways as key components of innate immune activation^29–32^. In CD4^+^ T cell subsets, *NFKB1* and *TRAF1* support involvement of TLR-NF-κB inflammatory signaling^33,34^, whereas in B cells, *IRF5* and *SLC15A4* further reinforce the importance of IFN/TLR coupling downstream of nucleic-acid sensing^35–37^. Additional B cell colocalizations at *BLK*, *CD83*, and *IL10RA* converge on pathways regulating BCR signaling, activation, and plasma cell differentiation^38–40^. Although the number of colocalized genes limits formal pathway enrichment analyses within each cell type, the repeated identification of genes with established roles in these immune pathways supports a model in which shared pathogenic processes in SLE are executed through distinct, lineage-specific regulatory programs.

In this study, sc-eQTLs were mapped across a multi-ancestry SLE cohort (102 AFR, 81 EUR, 62 SAS, 36 other ancestry) and tested for colocalization with a multi-ancestry SLE GWAS^11^ (5,877 EAS, 14,355 EUR, and 1,393 American SLE cases). Given the limited availability of large SLE GWAS datasets in SAS and AFR ancestries, this approach enabled testing of regulatory effects that may not be detectable in ancestry-stratified analyses alone, while also providing a broad survey of disease-relevant regulatory variation that is more generalizable across ancestries. In addition, differences in LD structure across populations, particularly the shorter LD blocks typically observed in AFR populations^41^, can serve as a powerful tool for refining putative causal variants even in the absence of dedicated fine-mapping, particularly where LD reference panels are less complete in understudied populations. However, LD heterogeneity across ancestries can also mean that a single causal variant exhibits distinct tagging patterns in different populations, potentially weakening the apparent concordance between GWAS and eQTL signals and reducing sensitivity to detect true colocalizations. Larger cohorts allowing stratified analyses will ultimately be required to clarify regulatory architecture with greater precision. More broadly, these findings highlight that adequately powered multi-ancestry studies, both GWAS and QTLs, will be key to resolving the full spectrum of genetic regulation in SLE, broadening the generalizability of discoveries, and potentially enabling the identification of ancestry-specific disease mechanisms.

A notable finding of this study is that many SLE-relevant colocalizations were not detectable in OneK1K^4^, a large healthy donor single-cell dataset. Several factors likely contribute to this, including differences in technical features and ancestry composition, but multiple observations support an impact of disease context. Several established SLE-associated genes, including *IRF7, ITGAM, IKZF3*^42^, showed substantially larger regulatory effects in SLE than in healthy controls, and at loci such as *FHL3*, distinct association architectures were displayed in the two cohorts. These findings are consistent with prior studies demonstrating that inflammatory environments can modify genetic regulatory effects^3^, potentially through cytokine-dependent chromatin remodeling and altered TF activity^43,44^. In autoimmune disease, where immune activation states are chronically perturbed, disease-specific eQTL effects may therefore represent an important but underexplored component of genetic risk architecture. Importantly, molecular disturbances in SLE precede clinical disease, with autoantibodies and altered interferon activity detectable years before clinical manifestations or classification^45,46^. Hence, the preclinical phase could provide a milieu in which SLE risk loci may contribute to disease development, with the regulatory effects persisting into the chronic disease state. Whether these disease-specific eQTL effects are a stable consequence of disease, driven by specific immune perturbations, or themselves causal to disease development remains an open question, and response eQTL experiments using SLE-relevant stimuli represent a promising avenue to begin addressing this.

Several limitations of this study should be considered. Although SLEmap is large relative to existing disease-focused single-cell eQTL maps, statistical power remains constrained, particularly for smaller cell populations and ancestry-stratified analyses. Furthermore, the limited availability of large ancestry-matched SLE GWAS datasets, particularly for African and South Asian ancestries, restricts ancestry-specific colocalization analyses, as disease-associated variants in these populations remain incompletely mapped. Future multi-ancestry SLE GWAS efforts will be essential to fully leverage SLEmap and uncover additional regulatory mechanisms. The comparison with OneK1K is also limited by differences in sequencing depth, cells per individual, and ancestry composition. SLEmap also included patients with variable disease duration and severity, making it difficult to distinguish genetic regulatory effects that contribute to disease from those that arise because of disease progression or activity. Larger studies spanning a broader range of longitudinal clinical states and incorporating sufficiently powered ancestry-matched healthy controls will help distinguish disease-specific regulatory effects from clinical, technical, and population-related variation.

Overall, SLEmap demonstrates that disease-specific, multi-ancestry single-cell regulatory atlases provide greater resolution for interpreting autoimmune disease genetics than healthy cohorts alone. By resolving the cellular and regulatory contexts through which SLE-associated variants act, this study advances understanding of how polygenic risk shapes immune dysfunction in SLE and provides a foundation for future mechanistic and therapeutic studies.

## Methods

### Sample Collection

Peripheral blood samples were collected from 300 female patients with SLE and 11 healthy controls. 213 SLE patients were recruited from Imperial Lupus Centre (Imperial College Healthcare NHS Trust, UK) and samples obtained from the Imperial College Healthcare Tissue and Biobank (ICHTB). ICHTB is approved by Wales REC3 to release human material for research (22/WA/0214). Ethical approval was provided by the ICHTB (Human Tissue Authority Licensing number 12275; Project number R14042-3A; sub-collection IMM_MB_13_001) and informed consent obtained from all participants. 87 SLE samples and 11 healthy control samples were collected from King’s College Hospital (King’s College Hospital NHS Foundation Trust, UK) and Guy’s Hospital (Guy’s and St Thomas’ NHS Foundation Trust, UK). Ethical approval was granted for this project under REC number 12/LO/1273, IRAS project ID 108060, and REC number 17/WA/0161 replaced by 22/WA/0214.

Inclusion criteria for SLE patients were female sex, age over 18 with prioritization of individuals under 50 years old, and clinician-diagnosed SLE according to the American College of Rheumatology (ACR) criteria with additional SLEDAI assessments. Recruitment prioritized individuals with European, African/Afro-Caribbean, and South Asian self-reported ethnicity. To minimize confounding, we prioritized recruitment of patients not treated with biologics within the past six months or with active infections requiring antimicrobials. Patients also had to be on a consistent, stable dose of their immunosuppressive or antimalarial medication for at least 12 weeks before the sample was taken. Patient demographics are summarized in **Supplementary Table 5**.

### WGS and scRNA-seq data generation

PBMCs were isolated from blood samples (collected into EDTA tubes) through density gradient centrifugation using Lymphoprep (STEMCELL Technologies, Canada) according to manufacturer instructions at each respective recruitment center. The cells were cryopreserved in FBS with 10% DMSO and stored in liquid nitrogen. For library preparation, PBMC samples were first thawed, washed with RPMI 1640 media containing 10% FBS, and rested for 16 hours at 37°C and 5% CO_2_. Upon harvesting, cells were resuspended in Cell Staining Buffer (BioLegend). Samples were counted and multiplexed into 69 pools, each containing four to five samples at equal cell density. To reduce background staining, cells were incubated with Human TruStain FcX (Biolegend) according to the manufacturer’s instructions. Surface proteins for CITE-seq were stained using TotalSeq™-C Human Universal Cocktail v1.0 (BioLegend), applied at a 1:6 dilution and targeting 137 surface proteins, including isotype controls. Following staining, cells were washed, stained with the live/dead dye 7-Aminoactinomycin D (7-AAD, BioLegend), and dead cells were removed by fluorescence-activated cell sorting (FACS).

Cells were processed using the 10X Chromium Next GEM Single Cell Immune Profiling 5’ v2 kit, as specified by the manufacturer’s instructions. 25,000 cells were loaded into each inlet of a 10X Chromium controller to create Gel Bead-in-emulsions (GEMs), with a targeted cell recovery of 12,000 cells per 10X reaction. Reverse transcription was performed on the emulsions, after which cDNA and CITE-seq supernatant were purified, amplified and used to construct RNA, CITE, T-cell receptor (TCR), and B-cell receptor (BCR) sequencing libraries. RNA, CITE, TCR, and BCR libraries were sequenced at a 10:1:1:1 ratio, respectively, using the Illumina NovaSeq6000 S4, with 100-bp paired-end reads.

For WGS, DNA was extracted from 10^6^ PBMCs remaining from single-cell sequencing (n = 291). For 20 individuals without any remaining PBMCs, additional whole blood samples were collected and used for DNA extraction. Extractions were performed using the automated QIAcube (Qiagen) and libraries were prepared using the NEBNext Ultra™ II DNA Library Prep Kit from Illumina. DNA libraries were sequenced in four batches over 19 lanes, with 15 to 16 individuals multiplexed at equimolar concentrations and sequenced per lane, aiming for 15x depth. DNA library pools were sequenced using the Illumina NovaSeq6000 S4 platform, with 150-bp paired-end reads.

### WGS data processing

Reads were aligned to the human reference genome (GRCh38.p13) using minimap2^47^ with default parameters for short reads. The resulting cram files were indexed and sorted using SAMtools^48^, and alignment quality metrics were assessed using Qualimap (v2.2.2)^49^ with duplicate reads marked using Picard. SNV calling was conducted with the Sarek pipeline (https://github.com/wtsi-hgi/sarek/), which incorporates haplotype calling and joint genotyping with GATK (version 4.4.0.0), following GATK best practices^50^. A total of 41,325,238 SNVs were called across 311 samples. All samples had genotype call rates over 97%.

Each sample was assessed for mapping quality, sex concordance, contamination, relatedness, variant missingness, and heterozygosity. Sex inference based on X/A depth and X/Y coverage ratios was concordant across all samples. Samples were filtered based on freemix score > 0.05 (evaluated using VerifyBamID2), PI_HAT > 0.1875 with more than 100 individuals (evaluated using PLINK), heterozygosity rate deviating by standard deviation from the cohort mean, and > 3% missingness, excluding 16 samples in total. Eight duplicate pairs were identified (PI_HAT > 0.98), and only the sample with lower contamination rate was used. One sample did not have matching single-cell data, and therefore excluded. Following QC, WGS data from 281 SLE patients and 5 healthy controls were carried forward for downstream analyses.

Variant QC was conducted to remove sequencing or mapping artefacts while retaining true genetic variants. Random forest classifiers were used following the approach described in Koko et al.^51^, using the WxS-QC pipeline^52^, which was adapted from the gnomAD QC framework (https://github.com/broadinstitute/gnomad_qc/). The truth set of variants used for training comprised high-confidence variants observed in 1,000 Genomes^53^, Omni 2.5 genotyping array (Illumina), 1,000 Genomes Indels present in Mills et al.^54^, and HapMap3 SNVs and Indels^55^. The negative set included variants meeting any of the following criteria: quality by depth <2, Fisher strand >60, or mapping quality <30. In total, 3,110,823 true-positive and 3,108,848 likely false-positive variants were used. A random forest (RF) classifier was then trained to distinguish true variants from artefacts using variant-level quality metrics as input features (**Supplementary Table 6**). Model performance was evaluated on chromosome 20 with SNVs ranked by RF score, the probability of being a true variant, and grouped into 100 bins. To define hard filters, combinations of RF score thresholds and genotype-level metrics were assessed.

The final filtering thresholds were set at RF score >86, genotype depth >5, genotype quality >20, and heterozygous allele balance >0.2. These criteria achieved a true positive rate of 99.5% and a false positive rate of 0.7%. Following quality control, 36,495,814 SNPs were retained for downstream analyses.

Genetic ancestry was inferred using KING^56^ (Kinship-based INference for Gwas) with the 1000 Genomes^53^ dataset as a reference. A sparse set of 17,491 “reliable” SNPs defined in AKT^57^ (ancestry and kinship toolkit) was used as input.

### Single-cell data processing

Raw single-cell data reads were processed and mapped with Cell Ranger Multi v7.0.0 (10X Genomics) using GENCODE release 32 (GRCh38.p13). QC steps were conducted as outlined in **Supplementary Fig. 14A**. For demultiplexing pools, YASCP (v.1.7) (https://github.com/wtsi-hgi/yascp) was used with genotyping information from the WGS data. Variants from the 1000 Genomes panel were used as a baseline set, supplemented by variants identified as differing between individuals within each pool. Among these, alleles expressed in the single-cell data were identified with cellSNP-lite (v1.2.3)^58^. Using this information, cells were assigned to each individual using Vireo (v0.2.3)^59^. Droplets classified as doublets or unassigned to individuals were excluded from further analysis. Five duplicate samples were also removed based on pairwise genetic correlation, and eight unidentifiable samples in pools with multiple samples without WGS data were excluded. This left 288 SLE samples and 10 healthy control samples for downstream analysis, of which 281 SLE samples and 5 healthy samples had matching WGS data (**Supplementary Fig. 14B**). Healthy control samples were retained in the cohort to facilitate robust annotation of canonical immune cell types.

Sequencing reads from the single-cell data were then split by individual using the vireo results and were used for personalized HLA mapping in order to improve the accuracy of HLA gene expression quantification. HLApm (https://github.com/davenportlab/HLApm) was used for this, by first generating a personalized HLA reference. HLA alleles for nine genes *HLA-A, HLA-B, HLA-C, HLA-DPA1, HLA-DPB1, HLA-DQA1, HLA-DQB1, HLA-DRA,* and *HLA-DRB1* were imputed at two-field resolution from WGS data using HLA*LA^60^, with the IMGT/HLA database (v3.42.0)^61^ as the reference. *HLA-DRB3, HLA-DRB4,* and *HLA-DRB5* alleles were inferred for each sample based on the known genetic architecture between these loci and specific *HLA-DRB1* alleles. Demultiplexed reads were then re-aligned to the personalized references using Cell Ranger as before and carried forward without additional droplet filtering.

The following QC steps were performed using YASCP with the exception of immune receptor processing. Low-quality droplets were removed if they contained fewer than 500 detected genes or had >20% mitochondrial reads. Gene expression complexity was scored by calculating the proportion of log_10_(number of genes expressed) to log_10_(total read counts) to evaluate the quality of gene expression data for each cell. Cells with scores less than 0.8 were removed. For each pool, droplets below 3 median absolute deviations (MAD) from the median for gene counts or read counts were removed, and droplets over 3 MADs from the median in mitochondrial read percentage were removed. Doublets were detected using multiple methods: scDblFinder^62^, scds^63^, DoubletFinder^64^, Scrublet^65^. Doublets detected with these methods largely agreed, and droplets were removed if they were detected using more than three methods. Immune receptor information was also used for droplet filtering. First, filtered BCR and TCR contigs from Cell Ranger were further processed using the Immcantation framework^66,67^ with V(D)J reassignment performed by IgBLAST^68^. Droplets containing more than one TRB chain, more than one BCR heavy chain, or both BCR and TCR chains were removed.

For normalization, read counts were scaled so that each cell had a total of 10,000 counts, and then logarithmic transformation was applied after a value of 1 was added to each count. From 36,600 genes sequenced, 2,000 highly variable genes (HVG) were identified using the Seurat V3 algorithm^69^.All immunoglobulin and T cell receptor related genes, including heavy and light chain V(D)J and constant region genes were excluded from HVG detection to prevent influence of clonotype, rather than cell type identity, on clustering. Multimodal integration between scRNA-seq and CITE-seq data was performed using totalVI (scvi-tools v1.1.2)^70^ with the HVGs and all 137 surface proteins.

UMAP generation was performed using neighborhoods constructed (k = 15) from 20 low-dimensional latent representations inferred by totalVI. The Leiden algorithm^71^ was used for unsupervised clustering with resolution 2. Two clusters with distinctively low ribosomal read percentage, as well as elevated mitochondrial read percentages and increased expression of nuclear-localized lncRNAs, such as *MALAT1*, appeared consistent with populations of ruptured or burst cells^72^ and were removed. Four clusters with abnormal CITE-seq profiles were also removed as they showed high levels of isotype control levels and co-expression of mutually exclusive cell surface markers which are indicative of non-specific antibody binding, a known phenomenon in CITE-seq experiments^73^. All removed clusters lacked consistent expression of known cell type-specific markers, and automated annotation using Azimuth (v0.5.0) with the Human PBMC reference^74^ failed to assign a uniform cell type to the cells within these clusters. Cells classified as erythrocytes or platelets from Azimuth were excluded from the dataset. The dataset was then re-integrated and re-clustered to obtain the final UMAP.

To manually annotate cell types, clusters were first broadly categorized as T cells, B cells, or “other” cells using surface protein markers and TCR/BCR data. This was followed by subsetting and reclustering within each group. Fine-level annotation was performed using canonical gene expression markers in combination with surface protein markers. Clusters displaying mixed or ambiguous marker expression were further reclustered to resolve distinct cell identities, leading to the annotation of 26 cell types. To complement manual annotation, automated annotation results from Azimuth^74^ were cross-referenced with manual labels to ensure accuracy.

### sc-eQTL mapping

SLE patients with matching single-cell and WGS data (n=281) were included in the sc-eQTL mapping. Genotypes for sc-eQTL mapping were derived from filtered WGS variants. Variants were retained if they had a MAF greater than 0.05, a missing genotype rate of 5% or less, and passed Hardy-Weinberg equilibrium (HWE) testing with a p-value threshold of 10^-6^. After applying these filters, a total of 6,152,421 variants were included in the analysis. Genotype PCs were then computed using these SNPs after LD pruning.

For gene expression, a mean pseudobulk approach was used to compute expression profiles per patient for each cell type, as described by Cuomo et al.^75^. Raw read counts were normalized to a target sum of 10,000 per cell to account for differences in sequencing depth. Counts were then log-transformed after 1 was added to each value to avoid taking the logarithm of zero. The mean expression was then computed for each cell type in each patient. For each cell type, only patients with at least 20 cells were included, and only cell types with at least 100 such patients were tested. All-PBMC expression profiles were generated by pseudobulking all cells from each donor, regardless of cell type.

Conditionally independent cis-eQTLs were mapped for each cell-type and for all-PBMC using tensorQTL^76^ within QTLight (https://github.com/wtsi-hgi/QTLight), testing SNPs over 5% MAF and within 1 Mb of the TSS. Associations between genotype dosage and pseudobulked gene expression levels were tested using multiple linear regression. To reduce variability and maximize statistical power to detect sc-eQTLs, only genes that were expressed in more than 5% of cells and in at least 10% of patients within each cell type were tested. The first 5 genotype PCs and an optimal number of gene expression PCs for each cell type were included in the model. The number of expression PCs to include was determined using a series of cis-eQTL scans using 0 to 50 expression PCs in increments of 5, and the number of expression PCs corresponding to the model where the number of significant eGenes plateaued was used (**Supplementary Table 7**). To determine statistical significance, 10,000 permutations were used to approximate beta-adjusted p-values, and genome-wide q-values were calculated using the Storey method, with a significance threshold of q-value < 0.05.

To understand which genes and SNPs are more likely to be detected in cell type-level eQTL mapping compared to all-PBMCs, the average gene expression level and cell type specificity of gene expression between eGenes in these groups were compared. To measure cell type specificity, Tau scores were calculated for each gene using the following formula.

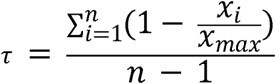

where *x_i_* is the expression in cell type *I,x_max_*, is the maximum value of (*x_1_,x_2_,…,x_n_*), and is the number of cell types. To assess differences between groups, a Kruskal-Wallis test was performed after confirming non-normality of the data using the Anderson-Darling normality test, and post hoc pairwise comparisons were conducted using the Wilcoxon rank sum test with Bonferroni multiple testing correction. With genes detected in both the cell type-level and all-PBMC analyses, eQTL effect sizes and the correlation between the effect sizes with eQTLs of identical lead SNPs were analyzed.

Enrichment of cell type-level eQTLs compared to all-PBMC eQTLs in distinct chromatin states were tested using the xGRviaGenomicAnno function from XGR^77^, applied to ChromHMM 15-state annotations^78^ derived from primary peripheral blood cell types in the Roadmap Epigenomics Project^79^. For this analysis, the genomic coordinates of all eSNPs were lifted over from hg38 to hg19, with 18,548 out of 18,608 eSNPs successfully converted.

For each tested cell type, ChromHMM annotations from matched primary immune cell types from peripheral blood were used. Statistical significance of enrichment was assessed at an FDR threshold of 0.05.

Pairwise sharing of sc-eQTLs across cell types was assessed using Multivariate Adaptive Shrinkage in R (mashr)^10^, which models effect sharing and heterogeneity by learning data-driven priors on the covariance structure of effect sizes, thereby accounting for differences in statistical power. For each pair of cell types, only genes tested in both were included (proportions shown in **Supplementary Fig. 15**). To estimate the covariance structure, a random subset of 200,000 gene-SNP pairs was used (random set), alongside a “strong set” comprising SNP-gene pairs that were significant in at least one cell type. For each cell type pair, the lead SNP (smallest P value) identified in one cell type (identification cell type) was evaluated in the other (replication cell type). Posterior effect sizes and local false sign rates (lfsr) were computed. An eQTL signal was considered shared if (i) the posterior effect sizes had the same direction, (ii) their magnitudes differed by less than twofold, and (iii) the lfsr was <0.05 in both cell types. The proportion of shared signals was defined relative to the number of significant signals in the identification cell type.

To formally assess signal sharing, the Flanders pipeline^80^ was used for fine-mapping and colocalization across eQTLs mapped in each cell type. Input parameters were set to a hole size (distance threshold for defining loci) of 200kb, a credible set threshold of 0.95, and was run with 1,000 max iterations. Flanders requires two significance thresholds to define genomic regions for fine-mapping. A primary significance threshold (p_thresh1) determines whether a region is selected for fine-mapping, for which the nominal p-value threshold from tensorQTL was used. The secondary inclusion threshold (p_thresh2) filters the set of variants included in the window and therefore determines the size of the fine-mapped region, which was determined using a scaled approach based on p_thresh1 with an upper bound to capture the full signal while constraining the region size:

p_thresh2 = min(10*p_thresh1, 1*10^-4^), if p_thresh1 < 1*10^-4^
p_thresh2 = min(10*p_thresh1, 1*10^-3^), if p_thresh1 < 1*10^-3^
p_thresh2 = min(5*p_thresh1, 1*10^-2^), if p_thresh1 < 1*10^-2^

Within Flanders, eQTL signals were fine-mapped using SuSiE^81^. Loci were excluded from fine-mapping if they exceeded the size threshold (n = 20) or did not yield any credible sets (n = 85). If an overlap in credible sets were found, colocalization was run using coloc^12^. Posterior probability for hypothesis 4 (PP.H4) > 0.8 were considered shared signals.

### Colocalization analysis with SLE GWAS

Colocalization analysis was conducted using coloc (v5.2.3)^12^ with summary statistics from a multi-ancestry SLE GWAS meta-analysis from Khunsriraksakul et al.^11^ Putatively independent GWAS loci for colocalization were defined using Locus Breaker (Sodbo Sharapov and Nicola Pirastu, Human Technopole; https://github.com/opentargets/gentropy). Briefly, input summary statistics were filtered at a p-value threshold of 1×10, and variants within 250,000 bp of one another were clumped together. Locus boundaries were subsequently defined as the positions of the leftmost and rightmost variants within each clump, extended by a flanking distance of 100,000 bp on each side. Loci overlapping the major histocompatibility complex (MHC) region (chr6:25,000,000–34,000,000; GRCh38) were excluded from all analyses to avoid spurious associations arising from complex LD in this region, leaving 197 independent GWAS loci for downstream testing. For genes harboring multiple conditionally independent eQTLs, nominal eQTL p-values were recalculated for each independent signal using a linear regression model in which all other independent SNPs at that locus were included as covariates, thereby isolating the association strength of each variant whilst accounting for the effects of other signals in the region.

For each GWAS locus, colocalization was only tested when at least 100 overlapping SNPs were present in the GWAS and sc-eQTL. coloc default prior probabilities were used (*p*1=1×10, *p*2=1×10 ^4^, *p*12 1×10^5^) for analysis. The coloc.abf() function was used to compute approximate Bayes factors and posterior probabilities. Colocalizations were defined as those with PP.H4 over 0.8. Colocalizations involving the 17q21.31 locus were excluded due to its known structural complexity^82^, including a megabase-scale inversion polymorphism that can result in reference-dependent mapping differences. For regional association plots, gene tracks were generated using LocusZoom^83^ with only coding genes displayed.

Novel colocalization events were identified using the Open Targets Platform^84^ (data release 25.12.0). All colocalizations with PP.H4 > 0.8 between credible sets from SLE GWAS studies (trait: MONDO_0007915) and eQTL or sc-eQTL studies were retrieved. To assess novelty, the lead variant from our analysis (the variant with the highest PP.H4) was compared with the lead variant from Open Targets (the variant with the highest posterior probability in the GWAS credible set). A colocalization was classified as known if the two lead variants fell within ±500 kb of one another and the associated eGene was the same; all remaining colocalizations were considered novel.

### Assessment of ancestry effects on colocalized eQTLs

To investigate the effect of ancestry on colocalized eQTLs, eQTLs were mapped separately within the three largest ancestry groups in the cohort (AFR, EUR, SAS) using the same eQTL mapping approach and multiple testing correction as the main analysis. Two genotype PCs and a number of expression PCs optimized for each cell type were included in the model (**Supplementary Table 7**). The same genes, individuals, and cell types as in the main eQTL mapping were used, with a minor allele frequency filter of >5% within each ancestry group. Effect sizes of colocalized eSNPs were compared across ancestry groups to assess concordance. LD for regional association plots was calculated within each ancestry group separately.

We tested the lead colocalized variants from those eQTL that colocalized with the GWAS for interactions with inferred genetic ancestry, accounting for genes with multiple independent eQTLs. We included only individuals with inferred AFR, EUR, and SAS ancestry, excluding those with EAS, AMR, or unassigned ancestry due to the small numbers of individuals in these groups. As this resulted in a subset of individuals compared to the original eQTL mapping, and inferred genetic ancestry was not included as a variable in the original eQTL mapping, we initially confirmed that the SNP-gene pair remained significant with inferred genetic ancestry added to the model, tested only in AFR, EUR, and SAS individuals. The following linear model was used to test ancestry interaction.

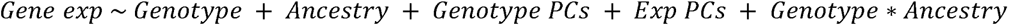

To prevent artefactual interactions from being detected due to nonlinear eQTL effects, we also compared the linear model to a model containing a second order polynomial of genotype and excluded those SNP-gene pairs where this nonlinear model fit better. SNPs were only tested if there were at least two reference and alternative allele homozygotes in each ancestry subgroup. Interaction p-values were calculated by comparing the model with no interaction to that with the interaction by ANOVA and corrected for multiple testing using the Benjamini-Hochberg method with significant interactions identified using an FDR threshold of 0.05. Interactions were plotted using the jtools R package^85^.

### Comparison with OneK1K

To assess whether SLEmap colocalizations could have been detected using a large publicly available healthy cohort, we compared our results against the OneK1K dataset. Single-cell data was obtained from CELLxGENE^86^. Only female individuals from OneK1K were included (n=565, 731,202 PBMCs) to align sex composition with SLEmap. Cells in the subsetted OneK1K dataset were clustered and visualized using the same method as SLEmap and manually reannotated using a shared set of RNA markers expressed in OneK1K to harmonize cell type labels across datasets. Average UMI per cell and average number of cells per individual were calculated from the female-subsetted OneK1K dataset across all cells.

Raw OneK1K genotype data underwent QC prior to imputation. Only biallelic SNPs were retained, and SNPs with missing rate >0.03, MAF <0.01, or deviation from Hardy-Weinberg equilibrium (p<10⁻³) were excluded. Individuals with missing genotyping rate >0.03, heterozygosity rate outside ±3 SD, or pairwise relatedness exceeding a GRM value of 0.125 were removed. The genotype data was then subset to the female individuals with matching scRNA-seq data (n=565) prior to imputation. Palindromic SNPs were removed and allele switches and strand flips corrected before imputation using the TOPMed Imputation Server^87,88^ with the TOPMed r3 reference panel^87^. Variants with imputation quality R2 > 0.8, MAF > 0.05, genotype missingness rate of 5% or less, and passing HWE testing with a p-value threshold of 10^-6^ were retained. This left 5,635,328 variants for eQTL mapping.

For eQTL mapping, identical expression and cell number filters were applied to OneK1K as in the main SLEmap analysis (genes expressed in >5% of cells and >10% of individuals, >20 cells per individual per cell type, >100 qualifying individuals per cell type). The same pseudobulk aggregation and eQTL mapping method was used with 4 genotype PCs and a cell type-specific number of gene expression PCs were used (**Supplementary Table 7**). Colocalization approach and priors were applied consistently across both datasets, using the SLE GWAS (Khunsriraksakul et al.) for colocalization. For each GWAS locus-eGene-cell type combination colocalized in SLEmap, we assessed whether the colocalization was replicated in OneK1K, excluding cell types not tested from OneK1K (cytotoxic CD4^+^ T cells and DN T cells). To isolate the contribution of disease status from ancestry, effect size comparisons with OneK1K were performed using eQTL mapping results from only European-ancestry individuals from SLEmap via mashR. Colocalizations with a posterior effect size ratio between SLEmap and OneK1K greater than 2 were classified as having a larger effect in SLEmap, while those with a ratio below 0.5 were classified as having a smaller effect in SLEmap.

### Functional characterization of colocalized loci

To investigate the functional basis of the cell-type-specific *TRAF1* colocalization in CM CD4^+^ T cells, we assessed whether the colocalized SNP disrupted TF binding motifs using motifbreakR^89^. Predicted disruptions were filtered to TFs expressed in the single-cell dataset, defined as expression in >5% of cells and >10% of individuals in at least one PBMC cell type. This retained PLAG1 as the sole candidate; TFAP2A and TFAP2C were excluded due to negligible expression in PBMCs, and TFAP2 was excluded as it represents a protein complex with no direct targets in the database. TF activity for PLAG1 was then scored in CM CD4^+^ T cells and memory B cells using decoupleR^90^ with the CollecTRI^91^ regulon database, and a two-sample t-test was used to compare activity between the two cell types.

Protein-protein interaction (PPI) analysis was performed using genes colocalized within the same cell lineage. Interactions were queried using the STRING database 13 (version 12.0, species = 9606), applying a high-confidence score threshold (≥700). Networks were examined to assess potential connectivity between colocalized genes.

## Acknowledgements

We thank all the patients for donating samples to this study. We thank the Wellcome Sanger Institute’s Scientific Operations team and Cytometry Core facility for generating the single cell RNA-seq and WGS data. We thank Steven Leonard, Matiss Ozols, Vivek Iyer and Wellcome Sanger Institute’s Human Genetics Informatics (HGI) team for single cell data analysis pipeline development. We also acknowledge the Open Targets partnership for supporting this work and for their valuable input. The Flanders pipeline was developed by the Biostatistics and Genome Analysis Units at Human Technopole, in particular thanks to Arianna Landini, Sodbo Sharapov, Edoardo Giacopuzzi, Bruno Ariano and Nicola Pirastu. We thank Angli Xue from Joseph Powell’s group (Garvan Institute of Medical Research) for sharing genotyping data from OneK1K. We acknowledge the support of the Imperial College Healthcare Tissue and Biobank, which is infrastructure funded by the NIHR Imperial Biomedical Research Centre. The views expressed are those of the authors and not necessarily those of the NIHR or the Department of Health and Social Care. For the purpose of Open Access, the author has applied a CC BY public copyright license to any Author Accepted Manuscript version arising from this submission.

## Author Contributions

Conceptualization: E.E.D., E.H., E.d.R., F.N., G.T., J.E.P., M.B., M.C.P., S.M., T.J.V.; Formal analysis: C.S., E.V.B., H.J., K.L.B., N.d.K., W.L.; Investigation: B.L.C., C.P.J., T.S.R.; Resources: C.W., J.E.P., M.B., M.C.P., M.W., N.B., T.J.V.; Original draft writing: C.S., H.J.; Review and editing: all authors; Supervision: C.P.J., E.E.D., G.T., J.E.P., T.J.V.; Funding Acquisition: E.E.D., E.d.R., F.N., G.T., J.E.P., M.B., M.C.P., T.J.V.

## Data availability

Raw single-cell data, raw WGS data, and genotypes called from the WGS data are available via European Genome-phenome Archive (EGAS00001008484). Processed single cell data and sc-eQTL summary statistics are available on Zenodo (10.5281/zenodo.21397133). SLE GWAS summary statistics were downloaded from https://liugroupstatgen.shinyapps.io/SLEv/. The Open Targets database is available at https://platform.opentargets.org/downloads.

OneK1K single cell data is available at CellxGene (https://cellxgene.cziscience.com/collections/436154da-bcf1-4130-9c8b-120ff9a888f2). Analysis code is available on Github (https://github.com/davenportlab/SLEmap).

