## Supplementary figures for "Single-cell multi-ancestry regulatory map of systemic lupus erythematosus"

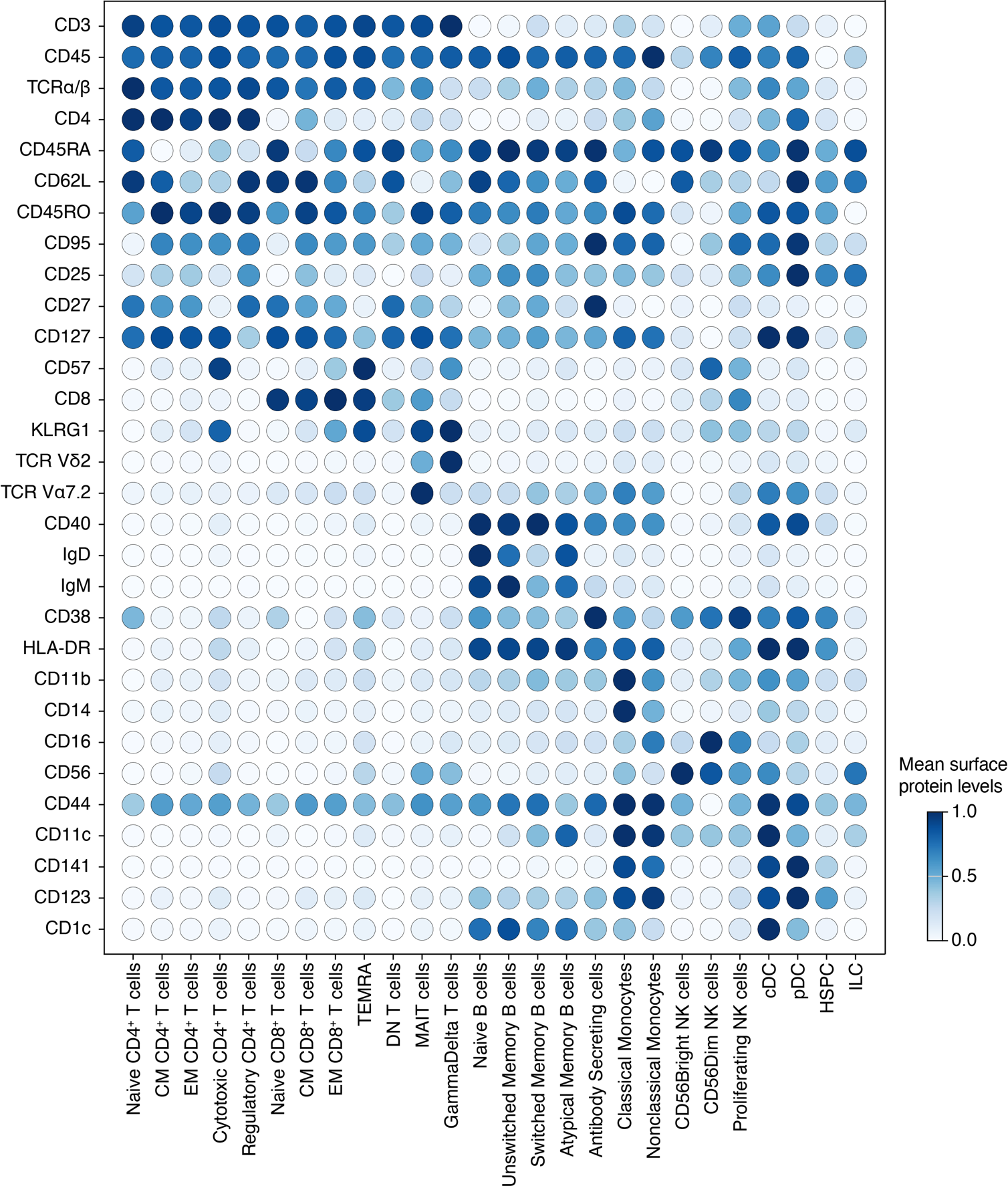

**Supplementary Figure 1.** Surface protein markers from CITE-seq data used to annotate immune cell subtypes. CITE-seq measurements were denoised and normalized using totalVI (Gayoso et al. 2021). Dot color indicates the average level of each surface protein within a cell type. CM=central memory; EM=effector memory; TEMRA=terminally differentiated effector memory RA cells; DN=double negative; MAIT=mucosal-associated invariant T; NK=natural killer; cDC=classical dendritic cell; pDC=plasmacytoid dendritic cells; HSPC=hematopoietic stem and progenitor cells; ILC=innate lymphoid cells

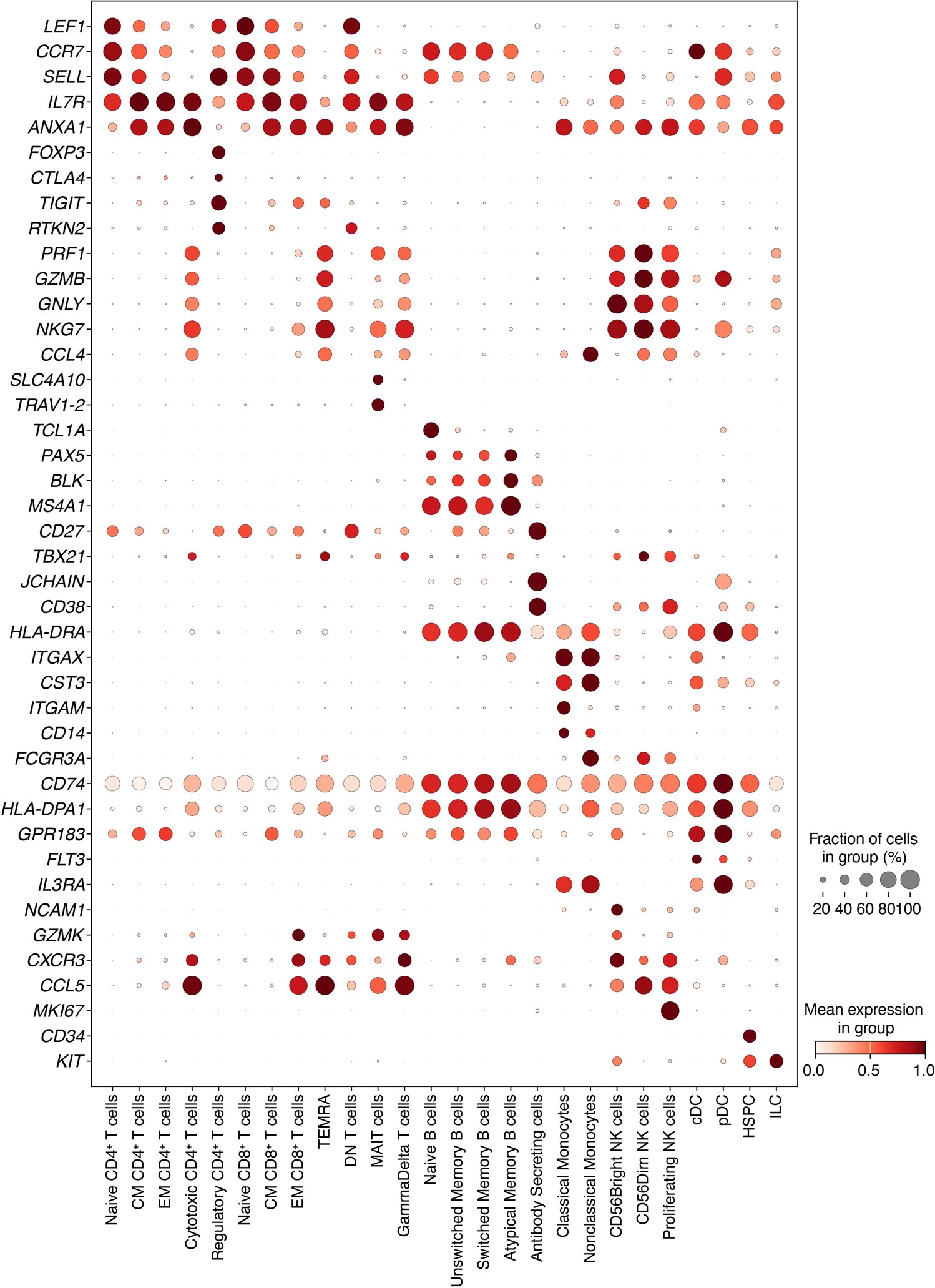

**Supplementary Figure 2.** RNA marker genes used to annotate immune cell subtypes. Expression values were normalized to 10,000 reads per cell and transformed using log(count+1). Dot color indicates the average expression level of each gene within a cell type, while dot size reflects the proportion of cells in which the gene is expressed.

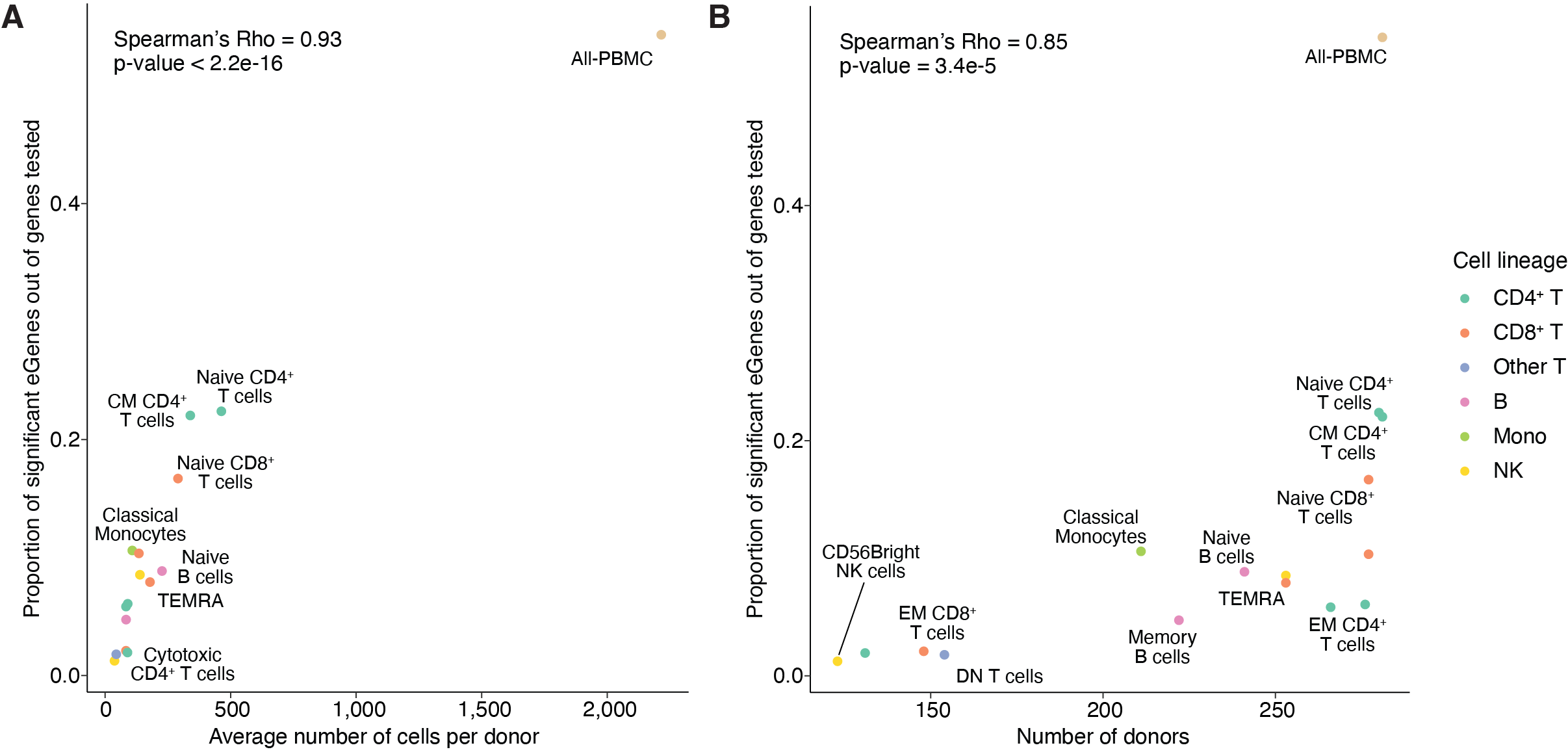

**Supplementary Figure 3.** Factors affecting the power to detect sc-eQTLs. **(A)** Correlation between the number of significant eGenes and the average number of cells per donor. Spearman's correlation rho was calculated to assess correlation. Cell types are colored by their cell lineage. Not all cell types are labelled due to space constraint. Details can be found in **Supplementary Table 2**. **(B)** Correlation between the number of significant eGenes and the number of donors included in the sc-eQTL mapping.

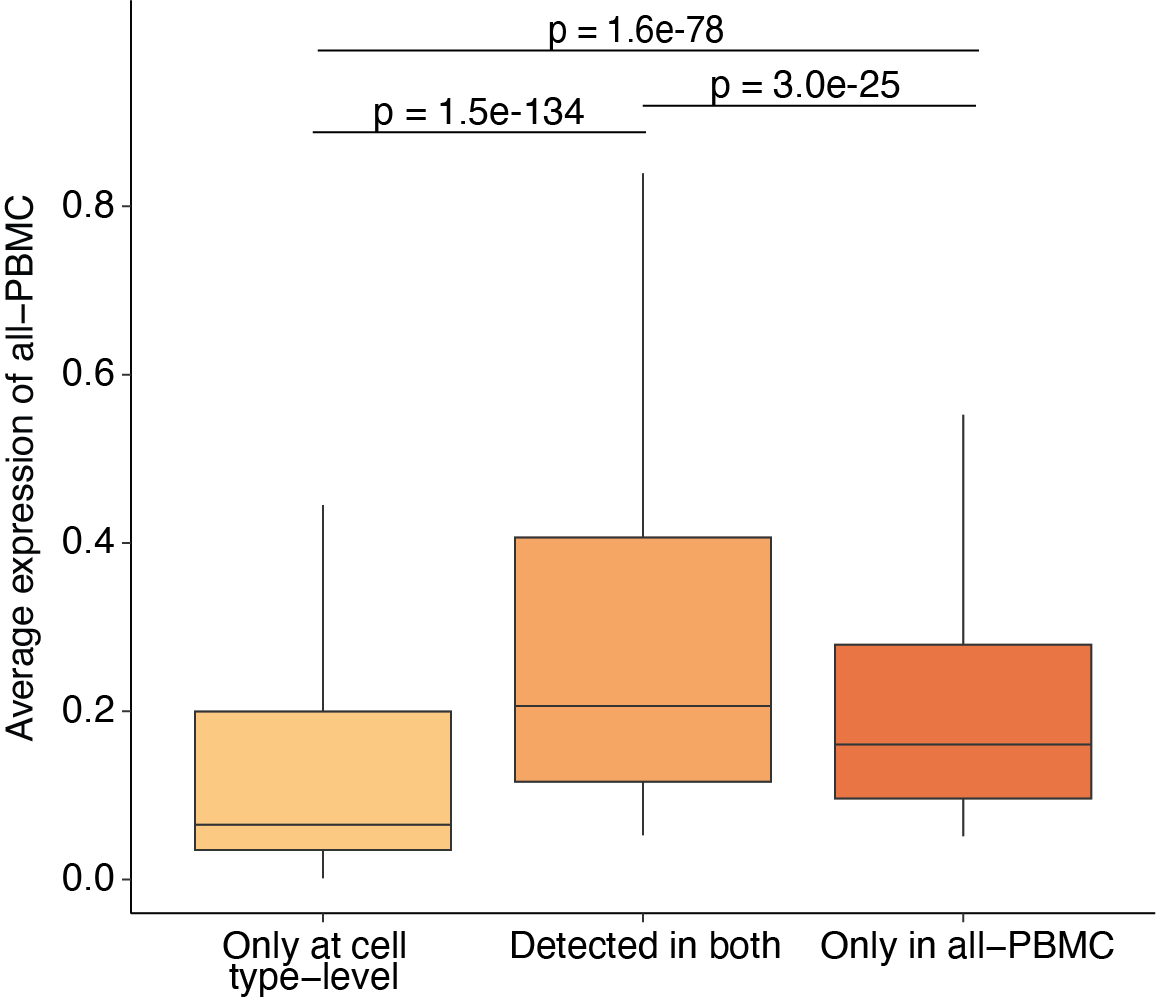

**Supplementary Figure 4.** Average expression level of genes in shared and distinct eGenes detected in all-PBMC eQTL mapping and cell type-level eQTL mapping. Differences between groups were tested using the Wilcoxon rank sum test.

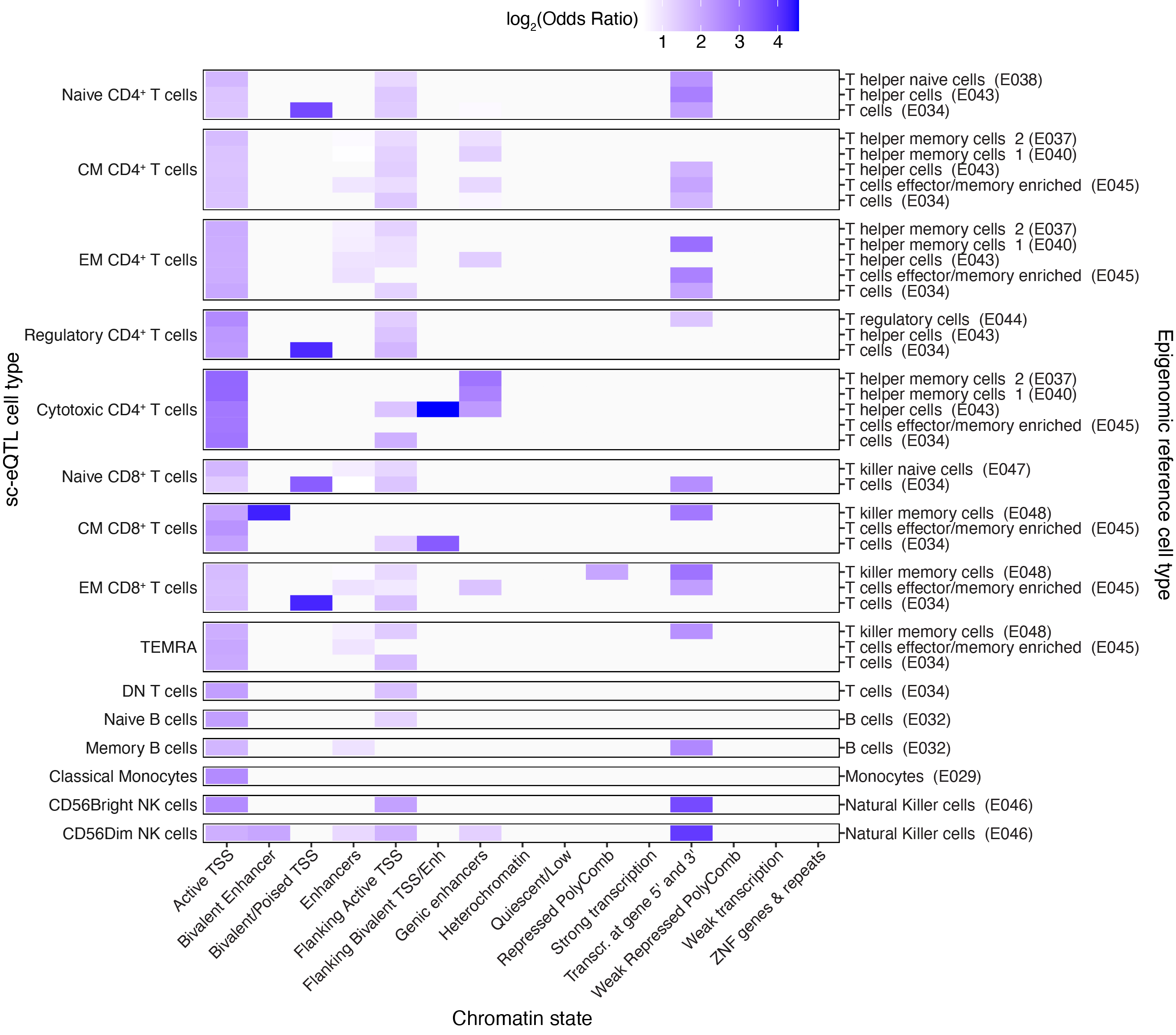

**Supplementary Figure 5.** Enrichment of cell type-level eQTL SNPs in chromatin states in contrast to all-PBMC eQTLs. Epigenomic annotations on matching primary peripheral blood cell types from ChromHMM were used. Only significant results are shown with color (FDR < 0.05). TSS=transcription start site; Enh=enhancer; ZNF=zinc finger

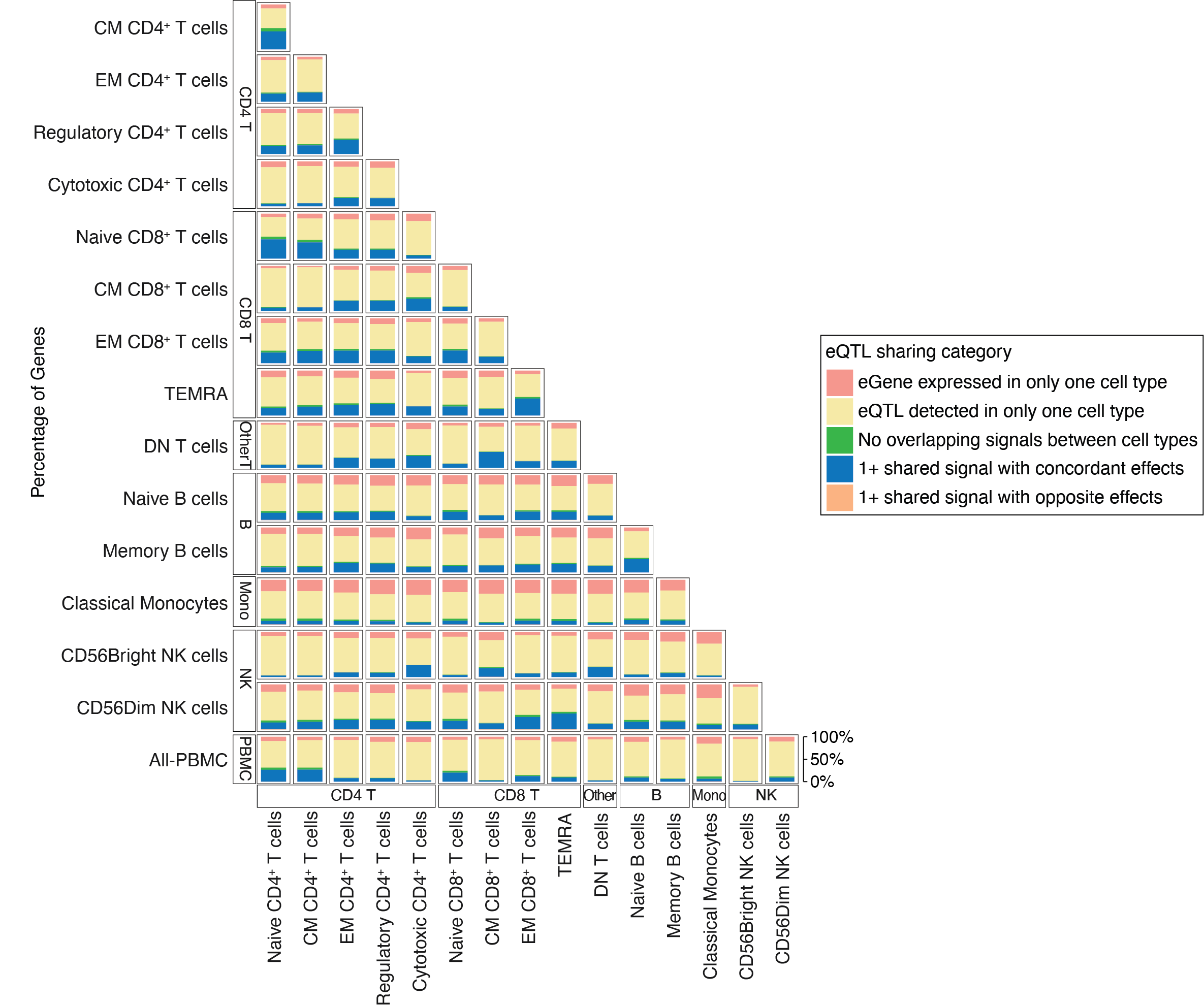

**Supplementary Figure 6.** eQTL sharing between cell types assessed using flanders by fine-mapping and colocalization. Shared signals were defined by colocalization posterior probability PP.H4 > 0.8. Opposite effects indicate that SNPs in both credible sets have effect sizes with opposite directions. A gene was considered expressed in a cell type if >10% of patients and >5% of cells had a count >1.

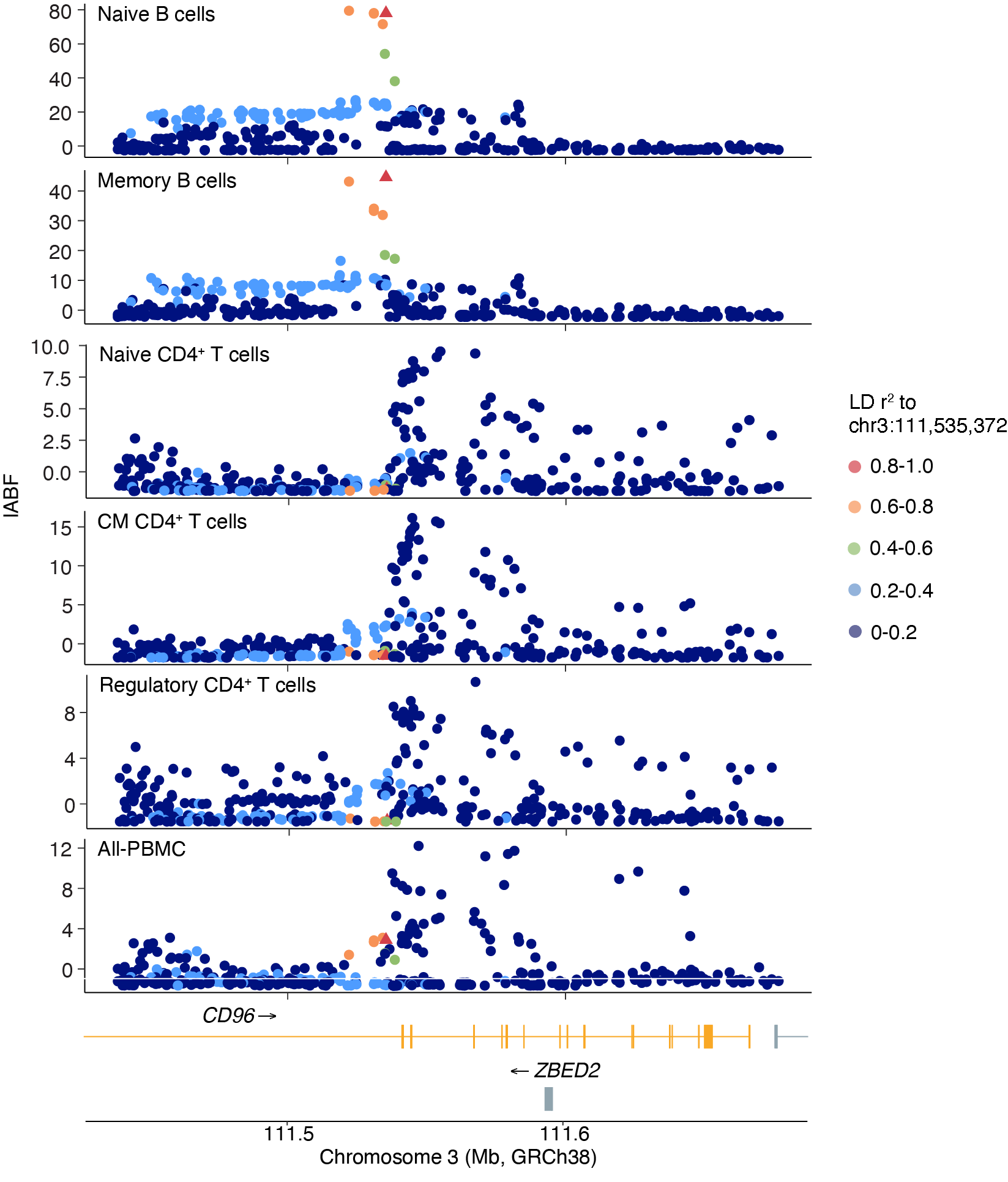

**Supplementary Figure 7.** Regional association plots of *CD96* sc-eQTLs across cell types. Naive and memory B cell signals colocalized with each other, with chr3:111,535,372 as the lead variant, while signals in naive, CM, and regulatory CD4^+^ T cells and all-PBMC colocalized with one another but not with the B cell signal. Points are colored by LD r² to chr3:111,535,372. lABF denotes the log approximate Bayes factor, where higher values indicate stronger evidence for association at each variant.

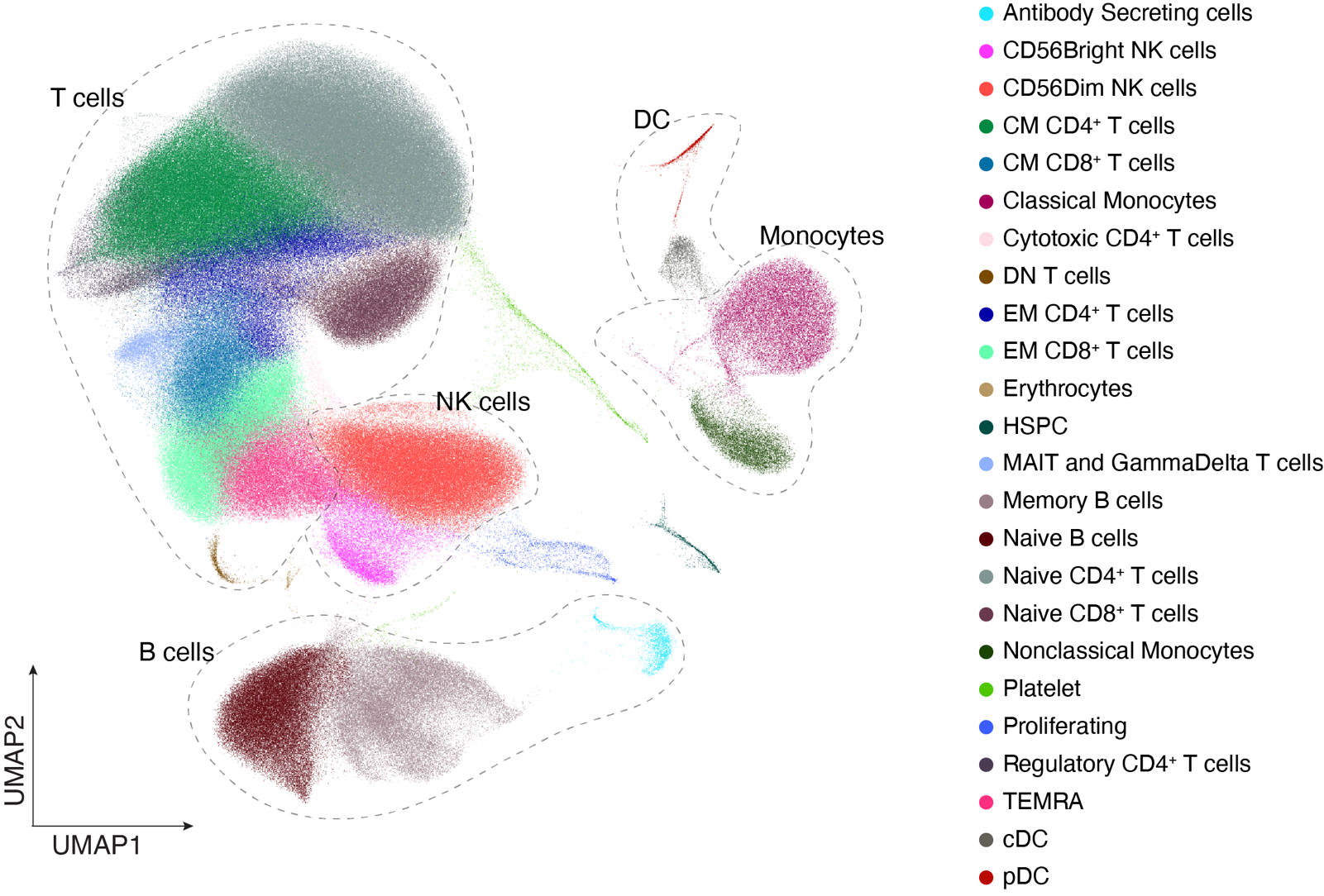

**Supplementary Figure 8.** UMAP of cells from female individuals from the OneK1K cohort colored by annotated cell type.

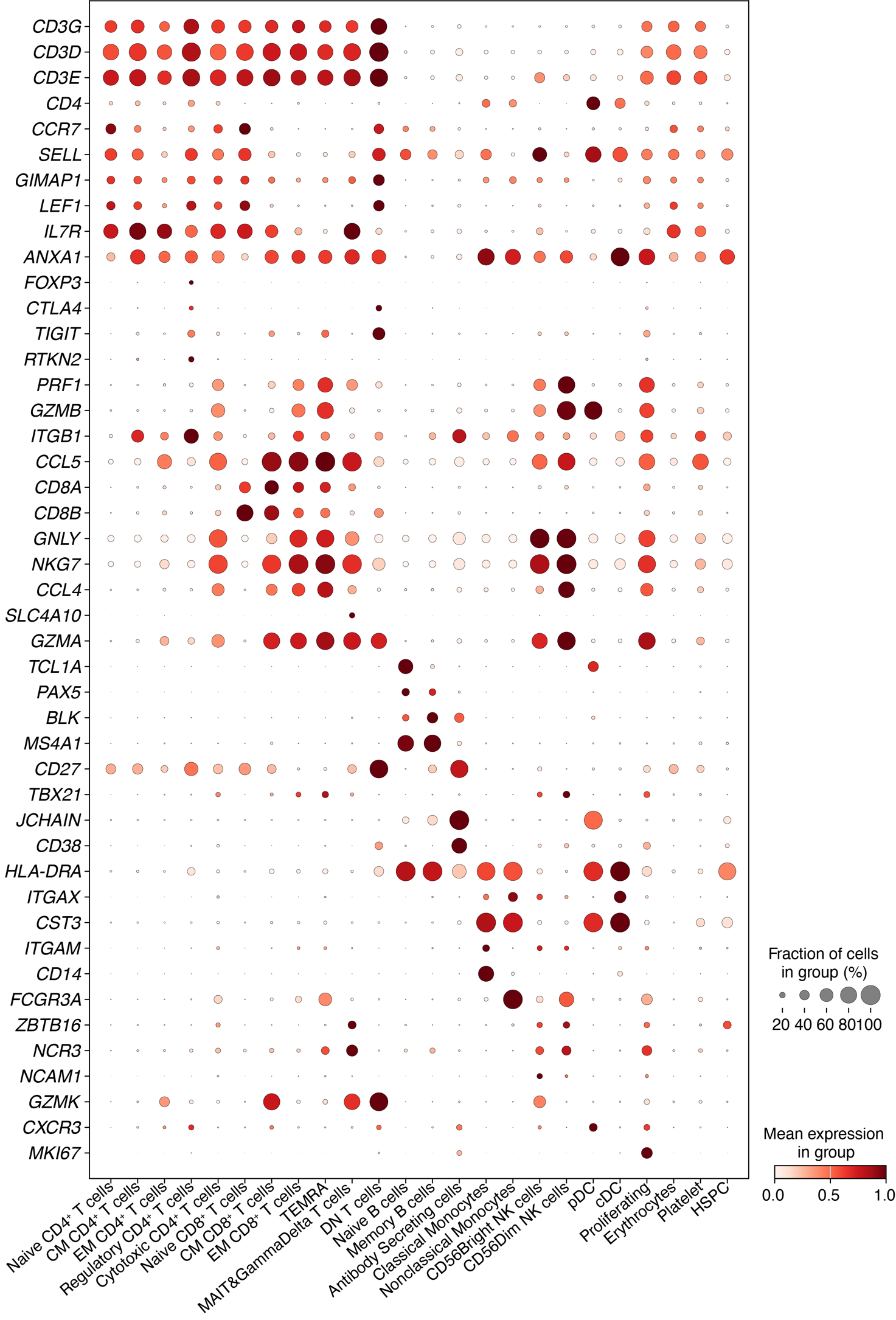

**Supplementary Figure 9.** RNA markers used for cell type annotation of OneK1K cells. Expression values were normalized to 10,000 reads per cell and transformed using log(count+1). Dot color indicates the average expression level of each gene within a cell type, while dot size reflects the proportion of cells in which the gene is expressed.

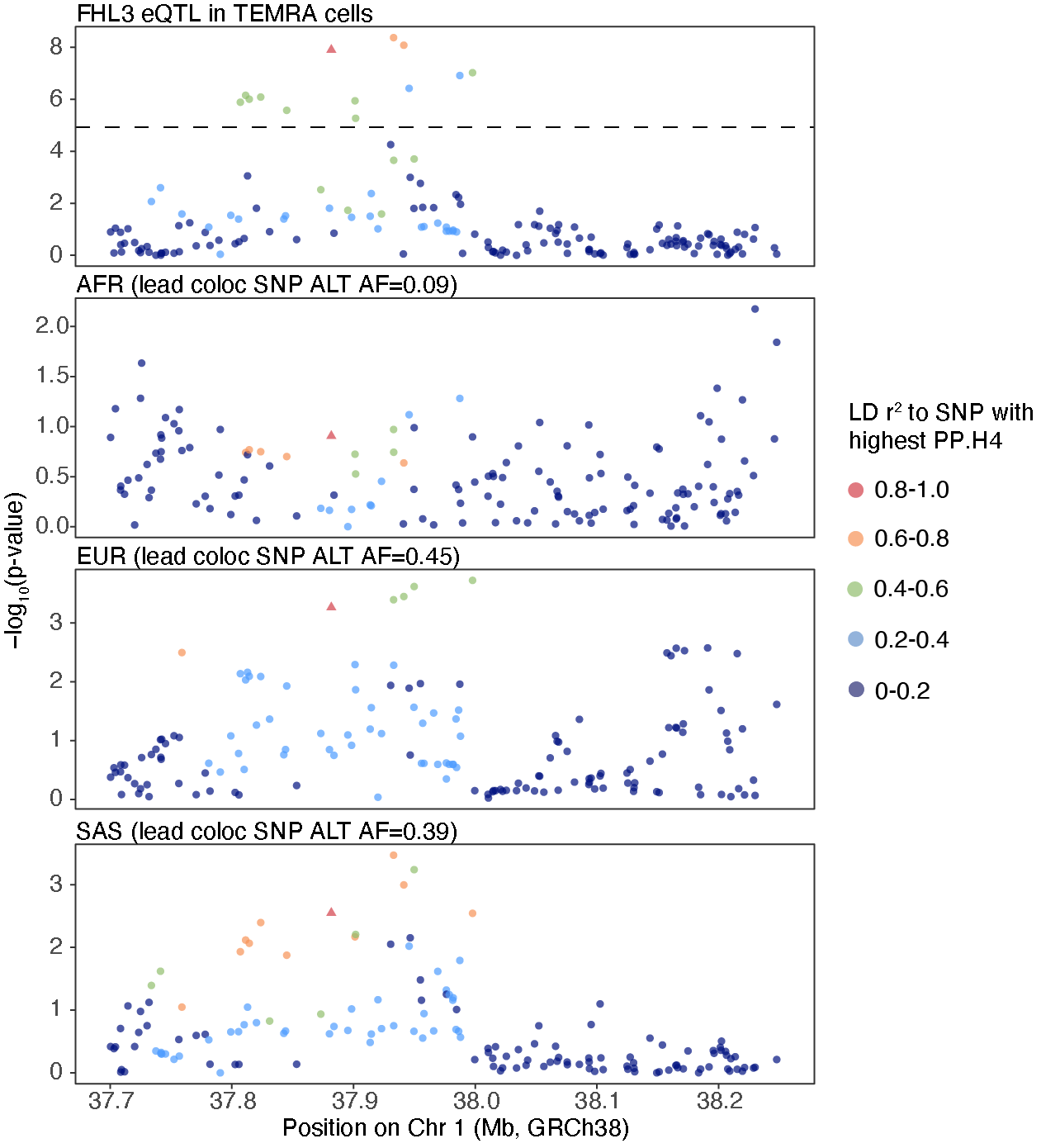

**Supplementary Figure 10.** *FHL3* eQTL in TEMRA by ancestry in SLEmap. Points are colored by LD r² to the colocalized SNV calculated within each ancestry group.

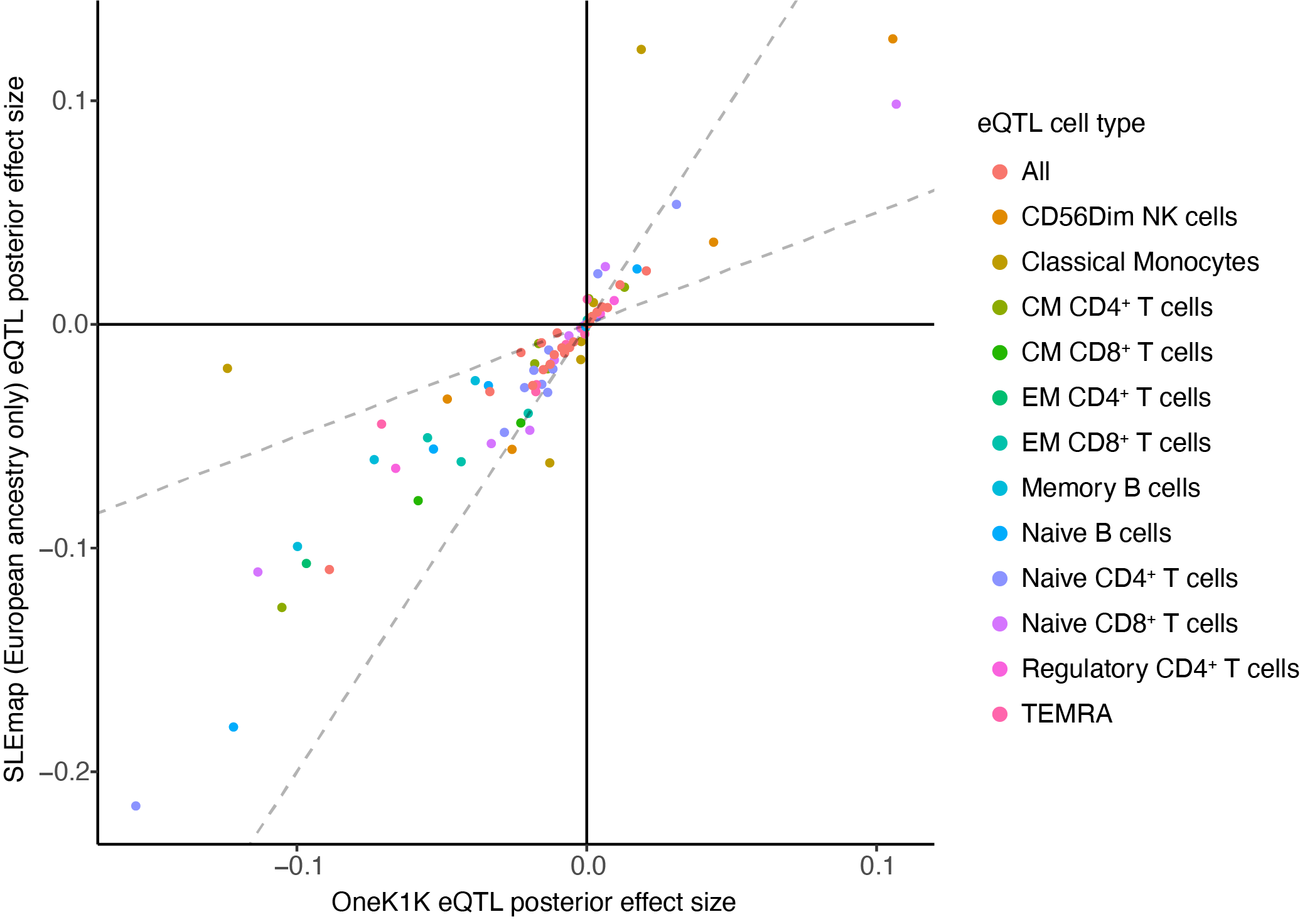

**Supplementary Figure 11.** Shared and distinct eQTL effect in colocalized signals between OneK1K and the European-only SLEmap subset colored by cell type. Posterior effect sizes calculated using MashR.

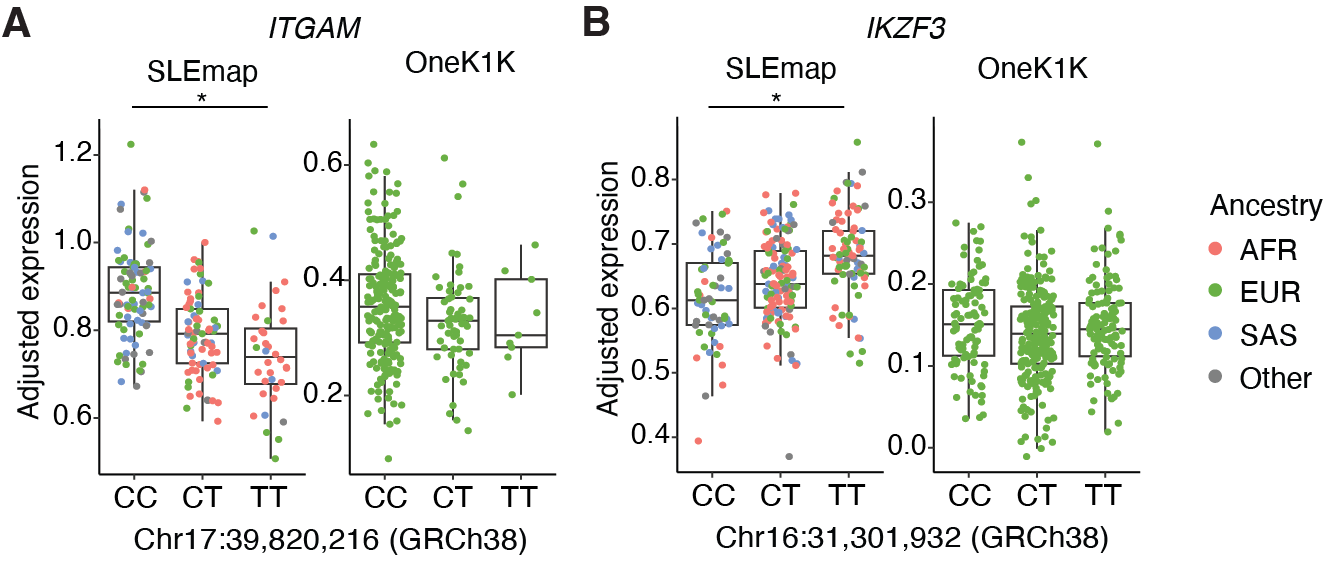

**Supplementary Figure 12.** Box plot of the *ITGAM* eQTL in classical monocytes and *IKZF3* eQTL in naive CD8^+^ T cells colored by ancestry.

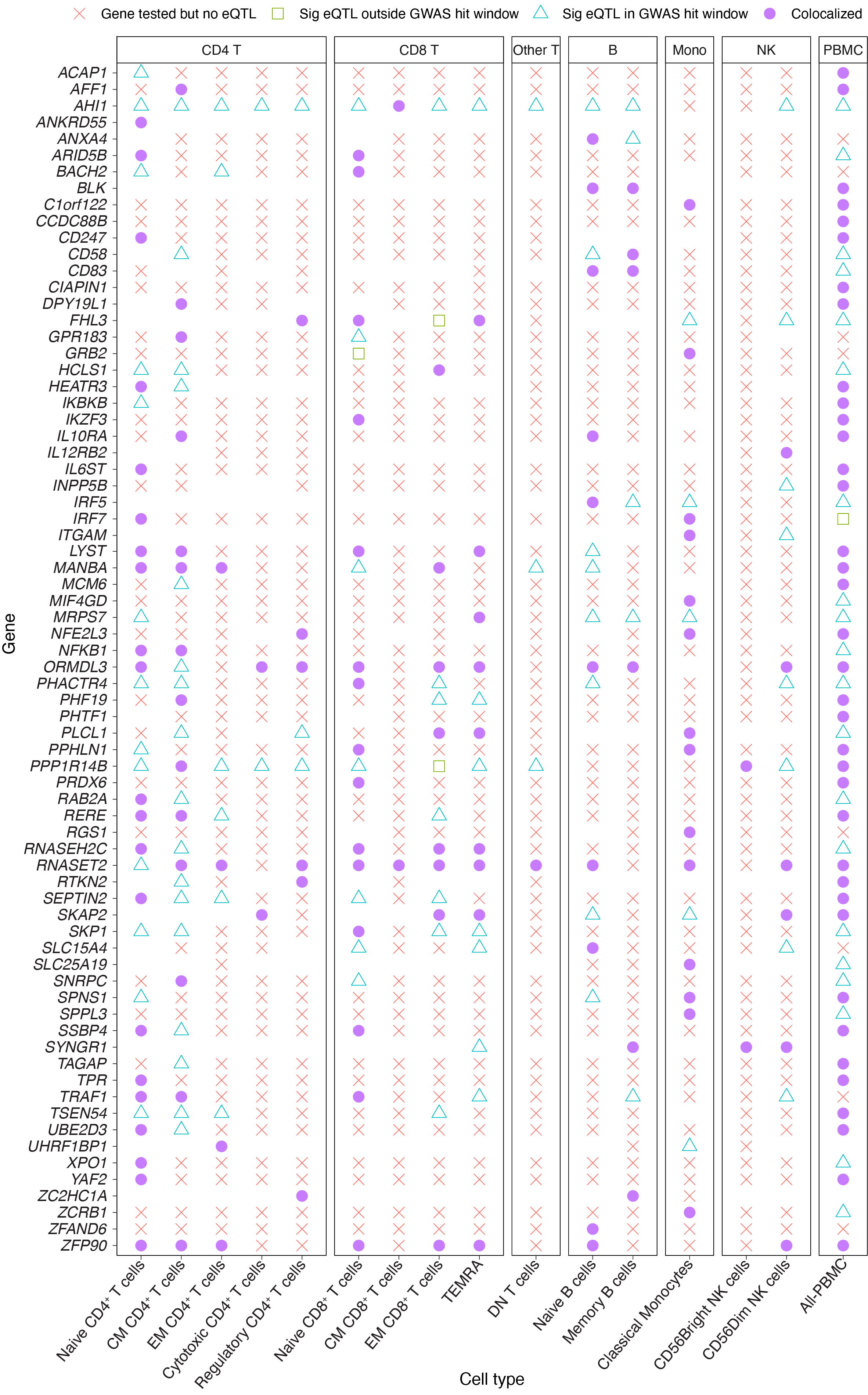

**Supplementary Figure 13.** sc-eQTL status for colocalized genes across cell types. Cell type-gene combinations with no symbol were not tested for eQTLs, due to insufficient expression (expressed in fewer than 5% of cells or fewer than 10% of donors). eQTLs were not tested for colocalization when fewer than 100 SNVs overlapped between the GWAS and the sc-eQTL dataset, or when the lead eQTL SNV lay outside the 1 Mb window around the lead GWAS SNV. For genes with multiple independent sc-eQTLs, the sc-eQTL with the higher likelihood of colocalization was plotted.

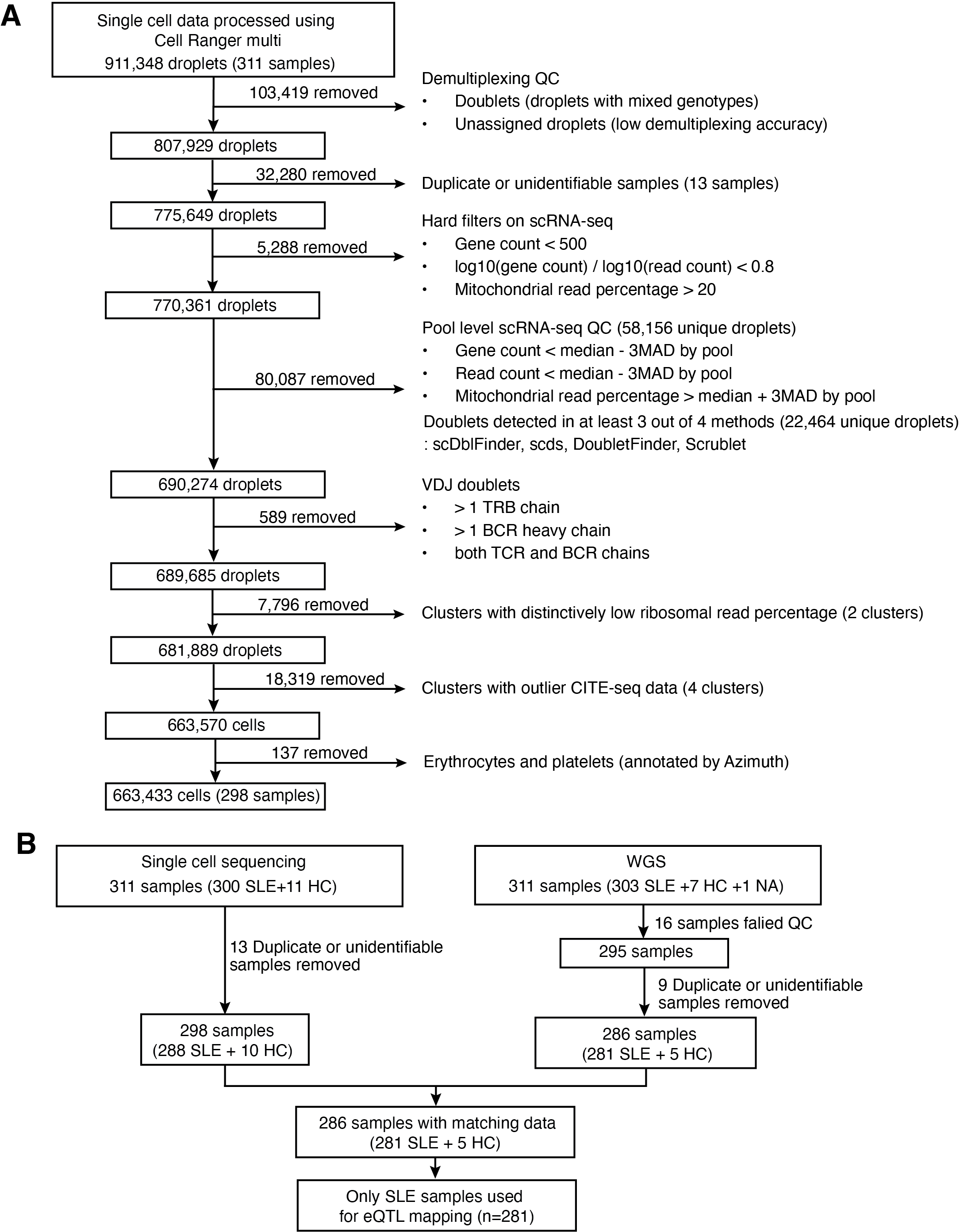

**Supplementary Figure 14.** Data QC flow charts. (**A**) Single-cell data QC steps to generate the dataset used for analysis. These steps were taken to filter out low-quality data and ensure reliable downstream analysis. MAD = median absolute deviation. (**B**) Sample-level QC flowchart for both single-cell data and WGS data.

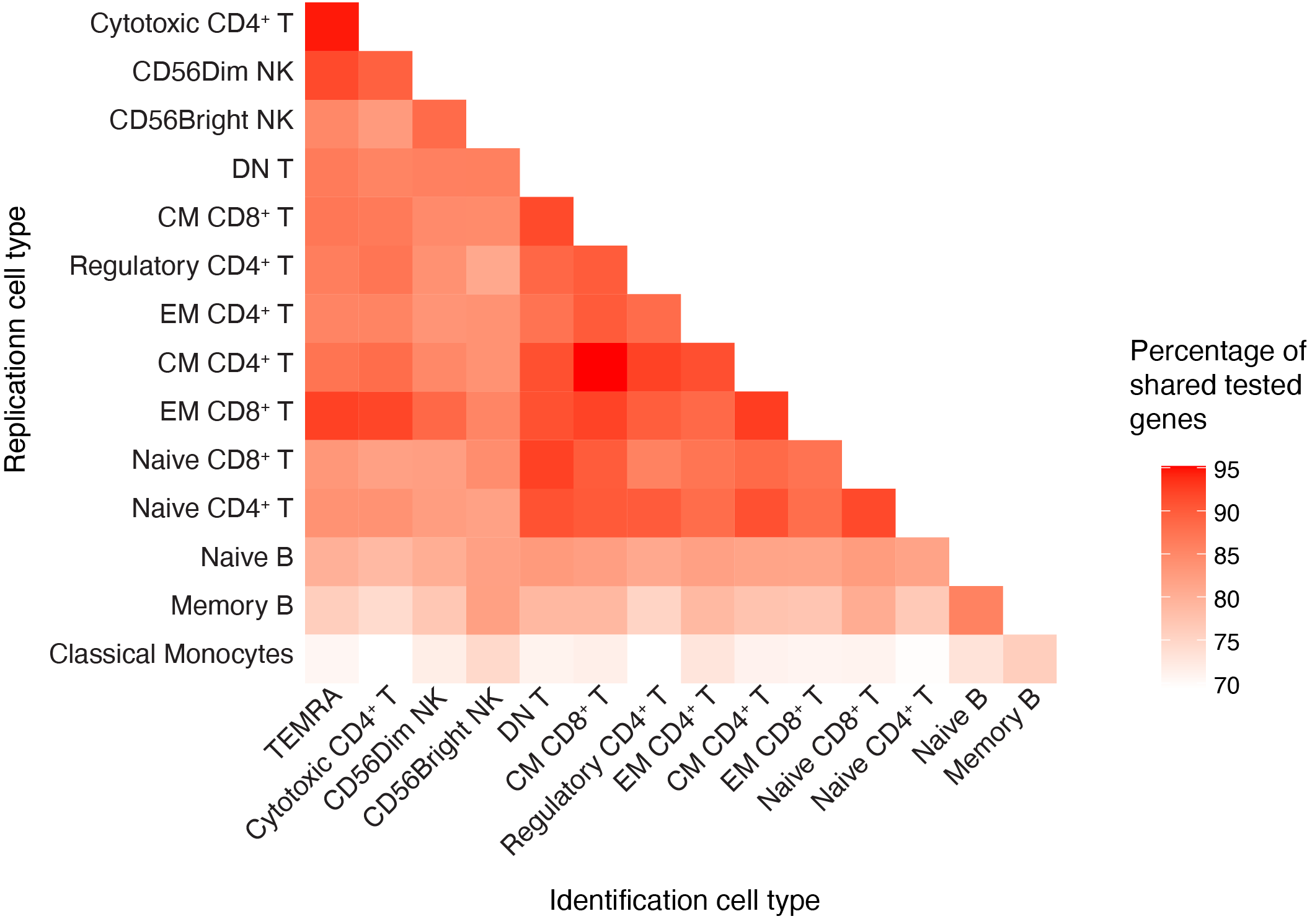

**Supplementary Figure 15.** Percentage of genes tested in both cell types for each pair, calculated relative to the total number of genes tested in either cell type. Cell types are ordered as in Figure 2D.
